# PIP_2_ electrostatically triggers vesicle fusion: arresting full SNARE assembly and vesicle fusion by PIP_2_-masking

**DOI:** 10.1101/2023.05.28.542642

**Authors:** Houda Yasmine Ali Moussa, Kyung Chul Shin, Janarthanan Ponraj, Sung Hyun Park, One-Sun Lee, Said Mansour, Yongsoo Park

## Abstract

SNARE proteins drive vesicle fusion and neurotransmitters release. Given that exocytosis is fast, and vesicle docking is tight, SNARE proteins are likely pre-assembled before fusion. However, the molecular mechanisms of the partially-assembled SNARE complex remain controversial. We use amperometry and the reconstitution of native vesicle fusion to show that MARCKS arrests basal fusion by masking PIP_2_ in a vesicle docking state where the SNARE complex is partially assembled. Ca^2+^/CaM or PKC-epsilon unmask PIP_2_ through the MARCKS dissociation, and thus rescue basal fusion and potentiates synaptotagmin-1-mediated Ca^2+^-dependent vesicle fusion. Our data provide the novel model that PIP_2_ electrostatically triggers vesicle fusion by lowering the hydration energy, and that masking PIP_2_ arrests vesicle fusion in a state of the partial SNARE assembly. Vesicle-mimicking liposomes fail to arrest vesicle fusion by masking PIP_2_, indicating that native vesicles are essential for the reconstitution of physiological vesicle fusion.

**One Sentence Summary:** Masking PIP_2_ by MARCKS arrests the full SNARE assembly and vesicle fusion.

## Introduction

Neurons and neuroendocrine cells communicate by neurotransmitter release and exocytosis^1^. Synaptic vesicles (SVs) store and release classical neurotransmitters such as glutamate, acetylcholine, GABA, and glycine, which mainly generate electrical signals to the postsynapse, whereas large dense-core vesicles (LDCVs) are responsible for exocytosis of amines, neuropeptides, and hormones, which mainly modulate synaptic activity^2–4^. Soluble *N*-ethylmaleimide-sensitive factor attachment protein receptor (SNARE) proteins mediate vesicle fusion^5, 6^ and synaptotagmin-1 is a Ca^2+^ sensor for fast Ca^2+^-dependent exocytosis of SVs and LDCVs^4^.

The time delay from presynapse to postsynapse for an electrical signal to travel across the synapse is ∼200 μs at 37 °C; this obervation suggests that ultrafast vesicle fusion might happen on the order of a few tenths of microseconds at physiological temperature after Ca^2+^ influx in the presynapse^7, 8^. In addition, SVs are tightly docked ∼1 nm from the active zone at synapses to be ready for fusion^9, 10^. Therefore, the SNARE complex seems to be partially assembled before fusion occurs: i) the SNARE motif is ∼ 7 nm long^11^; ii) the vesicle docking is tight, i.e., ∼ 1 nm distance between vesicle and the plasma membrane; and iii) vesicle fusion is fast on the order of a few tenths of microseconds to sub-milliseconds. However, the molecular mechanisms by which the partially assembled SNARE complex is stabilized and vesicle fusion is arrested before Ca^2+^ influx are unclear and under debate^4^.

Phosphatidylinositol 4,5-bisphosphate (PIP_2_) is a highly negative-charged lipid with −4 net charge at pH 7^12^, and is the second messenger in a variety of signaling pathways that regulate diverse cellular processes^13^. PIP_2_ has a critical function on vesicle fusion in neurons and neuroendocrine cells^14–16^. *In-vitro* reconstitution of vesicle fusion has shown that PIP_2_ is essential for triggering synaptotagmin-1-mediated Ca^2+^-dependent vesicle fusion by inducing the *trans*-interaction of synaptotagmin-1^17–20^. Here, we propose the novel model that PIP_2_ is also required for Ca^2+^-independent fusion by generating electrical double layer forces that lower the hydration energy barrier; masking PIP_2_ arrests vesicle fusion in a state of the partial SNARE assembly and tight docking.

Availability of free PIP_2_ is precisely and dynamically controlled by PIP_2_-binding proteins, and <u>m</u>yristoylated <u>a</u>lanine-rich <u>C</u>-<u>k</u>inase <u>s</u>ubstrate (MARCKS) is a well-characterized PIP_2_-binding protein^21^. MARCKS is highly conserved among vertebrates, in which it regulates neurodevelopment and synaptic functions^22, 23^. MARCKS is anchored by its N-terminus myristoylation in the plasma membrane, and the effector domain (ED, 25 amino acids) of MARCKS is enriched with 13 basic residues that electrostatically masks PIP_2_^24–26^. MARCKS can be dissociated from PIP_2_ by: i) Ca^2+^/Calmodulin (CaM), which disrupts the electrostatic interactions between MARCKS and PIP_2_^27, 28^; and ii) protein kinase C (PKC), which phosphorylates the ED, and thereby causes dissociation of MARCKS, like an electrostatic switch^25, 29^.

Synaptotagmin-1 is the primary Ca^2+^ sensor for fast exocytosis by inducing local membrane bending that lowers energy barrier for fusion^20^. Intriguingly, CaM is considered as an additional Ca^2+^ sensor for vesicle fusion^30^; e.g., vacuoles in yeast^31^, endosomes^32^, cortical granules in sea urchin eggs^33^, LDCVs^34–37^, and SVs^38–41^, although the contributing function of Ca^2+^/CaM on vesicle fusion as a Ca^2+^ sensor is controversial. The mechanisms how Ca^2+^/CaM and PKC regulate both Ca^2+^-dependent and Ca^2+^-independent vesicle fusion remain poorly understood.

By using the interdisciplinary approaches including amperometry to monitor exocytosis in real time and the *in-vitro* reconstitution of native vesicle fusion in a physiological context, we demonstrate that MARCKS arrests Ca^2+^-independent basal fusion by masking PIP_2_; SNARE complex is partially assembled in a state of vesicle docking. Ca^2+^/CaM unmasks PIP_2_ by dissociating MARCKS, and thus rescues basal fusion and potentiates synaptotagmin-1-mediated Ca^2+^-dependent vesicle fusion. For Ca^2+^-independent fusion, activation of PKC-epsilon triggers basal fusion of LDCVs by unmasking PIP_2_. Importantly, vesicle-mimicking liposomes (V-liposomes) fail to reproduce the PIP_2_ effect on basal fusion, and therefore purified native vesicles, i.e., LDCVs and SVs, are essential for the reconstitution of physiological vesicle fusion; this result is consistent with a previous report on the cholesterol effect^20^.

## Results

### Requirement of PIP_2_ for Ca^2+^-independent basal fusion

We have established the reconstitution system of vesicle fusion *in-vitro* by using purified native vesicles such as LDCVs and SVs^17, 18, 20, 42–44^. Native vesicles readily fuse with the plasma membrane-mimicking liposomes (PM-liposomes) that contain the stabilized Q-SNARE complex of syntaxin-1A and SNAP-25A in a 1:1 ratio by a C-terminal fragment of VAMP-2^45^ (**Fig. 1**). The depletion of PIP_2_ results in dramatic disruption of LDCV exocytosis in chromaffin cells^14^, so we first examined the effect of PIP_2_ on basal fusion of LDCV by using the reconstitution system. Intriguingly, Ca^2+^-independent basal fusion of LDCV was not observed when PIP_2_ was excluded from PM-liposomes prepared by the direct method; large unilamellar vesicles (LUVs) with ∼110 nm in diameter (**Online Methods** for detail). Basal LDCV fusion without Ca^2+^ was robust when PM-liposomes contained 1% PIP_2_ (**Fig. 1a**). Pre-incubation of VAMP-2_1-96_ (i.e., the soluble cytoplasmic region of VAMP-2) with PM-liposomes completely blocked vesicle fusion by occupying the free Q-SNARE complex; this result confirms that vesicle fusion is SNARE-dependent. Absence of PIP_2_ in PM-liposomes led to complete inhibition of basal LDCV fusion, comparable to the result of VAMP-2_1-96_ treatment (**Fig. 1a**).

**Figure 1.**
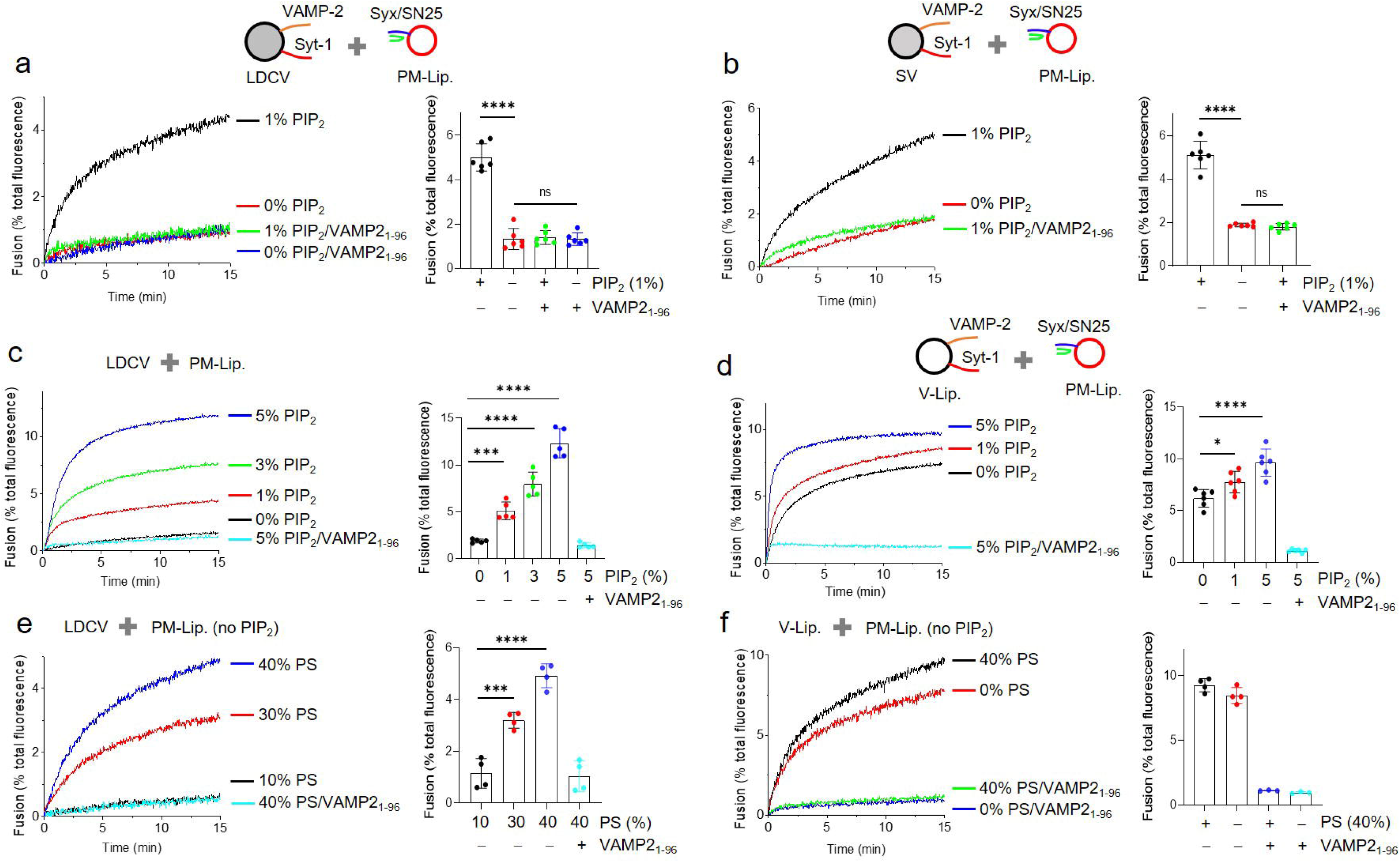
PIP_2_ is required for Ca^2+^-independent basal fusion. *In-vitro* reconstitution of vesicle fusion using a fluorescence dequenching-based lipid-mixing assay. (**a,b**) No fusion happens when PIP_2_ is absent in PM-liposomes. Purified LDCVs (**a**) or SVs (**b**) were incubated with PM-liposomes that incorporate the stabilized Q-SNARE complex of syntaxin-1A/SNAP-25A (Syx/SN25) in a 1:1 molar ratio; PM-lip., large unilamellar vesicles (LUVs) prepared by the direct method. PM-liposomes contain either 0% or 1% PIP_2_. (**c**) Effect of PIP_2_ dose on basal fusion of LDCV. (**d**) Instead of native LDCVs and SVs, V-liposomes (V-lip.) that incorporate full-length synaptotagmin-1 and VAMP-2 were incubated with PM-liposomes. Effect of PIP_2_ dose on basal fusion of V-lip. with PM-lip. (**e,f**) PIP_2_ is not included in PM-liposomes. Effect of PS dose on basal fusion of LDCV (**e**) or V-lip. (**f**) with PM-lip. Lipid composition of PM-liposomes in **a-d**: 45% PC, 12% PE, 3% labeled PE, 10% PS, 25% Chol, 4% PI, and 1% PIP_2_. When PIP_2_ concentration was changed, PI contents were adjusted accordingly. Lipid composition of PM-liposomes in **e,f**: 20% PC, 12% PE, 3% labeled PE, 25% Chol, and 40% PS. When PS contents were changed, PC contents were adjusted accordingly. Lipid composition of V-liposomes: 55% PC, 20% PE, 15% PS, and 10% Chol. Physiological ionic strength with 1 mM MgCl_2_/3 mM ATP was used in all experiments; ATP removes contaminating Ca^2+^ as a Ca^2+^ chelator^19^. Preincubation of PM-liposomes with VAMP-2_1-96_, the cytoplasmic domain of VAMP-2 (aa 1-96), specifically disrupted SNARE-dependent vesicle fusion. Data are means ± SD from four to six independent experiments. One-way ANOVA test with Bonferroni correction was used in **a-c,e,f**. One-way ANOVA with post-hoc Turkey test was applied in **d**; *, *p* < 0.05, ***, *p* < 0.001, ****, *p* < 0.0001.

Next, we tested the PIP_2_ effect on basal fusion by using SVs isolated from mice brains (**Fig. 1b**). Indeed, absence of PIP_2_ in PM-liposomes (0% PIP_2_) led to no SV fusion; again, SV fusion was fully disrupted to the level of VAMP-2_1-96_ treatment (**Fig. 1b**), similar to the results of LDCV fusion. LDCV fusion was completely blocked when PM-liposomes contained no PIP_2_, and increasing PIP_2_ concentration from 1% to 5% significantly increased LDCV basal fusion in a dose-dependent manner (**Fig. 1c**). Due to lipid/protein composition and the structural integrity of native vesicles, they have advantages for a fusion assay to reproduce endogenous vesicle fusion in a physiological context^17, 20^. To confirm that Ca^2+^-independent basal fusion depends on PIP_2_, vesicle-mimicking liposomes (V-liposomes) were used instead of purified native vesicles. V-liposomes incorporated full-length VAMP-2 and synaptotagmin-1. In contrast to native vesicles, V-liposome fusion still occurred even in the absence of PIP_2_ in PM-liposomes (0% PIP_2_), but increasing PIP_2_ concentration slightly increased liposome-liposome fusion (**Fig. 1d**). Taken together, these results indicate that PIP_2_ is required for Ca^2+^-independent basal fusion of native vesicles (i.e., LDCV and SV), but is not essential for liposome-liposome fusion, although PIP_2_ slightly increases the efficiency of liposome-liposome basal fusion.

The methods of liposome preparation might affect fusion efficiency and the PIP_2_ effect. The direct method of liposome preparation is to incorporate proteins in pre-formed liposomes, whereas the standard method uses co-solubilizing lipids and proteins, then proteoliposomes form after detergent removal (**Online Methods** for detail). We therefore assessed PM-liposomes prepared by the standard method, which are small unilamellar vesicles (SUVs) with diameter ∼50 nm (**Supplementary Fig. 1**). As expected, Ca^2+^-independent basal fusion of LDCV (**Supplementary Fig. 1a**) and SV (**Supplementary Fig. 1b**) was completely blocked when PM-liposomes (SUV) contained no PIP_2_ (0% PIP_2_). We also confirmed that LDCV fusion did not occur without PIP_2_ in PM-liposomes when the full-length syntaxin-1A and SNAP-25A binary acceptor complexes (**Online Methods** for detail) were included in PM-liposomes (**Supplementary Fig. 1c**). Altogether, the requirement of PIP_2_ for Ca^2+^-independent basal fusion of native vesicles is reproduced regardless of different liposome preparation methods and the full-length SNARE complex.

### Anionic phospholipids are required for basal fusion

PIP_2_ has a net charge of −4 at pH 7^12^, and is among the most highly negative-charged lipids. To test the hypothesis that the PIP_2_ dependence for Ca^2+^-independent basal fusion is mediated by the electrostatic effect, we replaced PIP_2_ with phosphatidylserine (PS), which has a net charge of −1 at neutral pH^12^. LDCV basal fusion was not observed when PM-liposomes contained < 10% PS, but increasing PS contents (to 30∼40%) triggered basal fusion (**Fig. 1e**); LDCV fusion with PM-liposomes containing 40% PS was equivalent to that of PM-liposomes that contained 1% PIP_2_. Similar to PIP_2_-independent liposome-liposome fusion (**Fig. 1d**), V-liposome fusion was robust even without PS in PM-liposomes (**Fig. 1f**). These results suggest that anionic phospholipids evoke basal fusion of native vesicles by the electrostatic effect. Basal fusion of liposome-liposome occurs regardless of anionic lipids; PIP_2_ can increase liposome-liposome fusion (**Fig. 1d**).

### SNARE assembly and the ternary SNARE complex formation despite no basal fusion

Next, we investigated whether SNARE proteins of SVs were assembled in the absence of PIP_2_ in PM-liposomes. SNARE assembly was monitored using anisotropy measurement (**Fig. 2a**). The VAMP-2_49-96_ fragment labeled with Alexa Fluor 488 (Alexa 488-labeled VAMP-2_49-96_) is incorporated in the stabilized Q-SNARE complex in PM-liposomes, and then the Alexa 488-labeled VAMP-2_49-96_ fragment is displaced from the stabilized Q-SNARE complex as the full-length VAMP-2 zippers in the N-to-C terminal direction^45^. The displacement of the Alexa 488-labeled VAMP-2_49-96_ fragment by endogenous VAMP-2 of native vesicles leads to a decrease in fluorescence anisotropy because of the increased rotational mobility of fluorophore^17, 42^; this result shows that the displacement of Alexa 488-labeled VAMP-2_49-96_ is a reporter of SNARE assembly. We observed no difference in SNARE assembly and kinetics of the Alexa 488-labeled VAMP-2_49-96_ displacement when SVs are engaged in SNARE assembly with PM-liposomes containing either 0% or 1% PIP_2_ (**Fig. 2a**), although SV fusion was completely arrested without PIP_2_ in PM-liposomes (**Fig. 1b**). As a control, the pre-incubation of SVs with the soluble and unlabeled Q-SNARE complex occupied free endogenous VAMP-2, so anisotropy did not change significantly, confirming the engagement of endogenous vesicular VAMP-2 in SNARE assembly (**Fig. 2a**).

**Figure 2.**
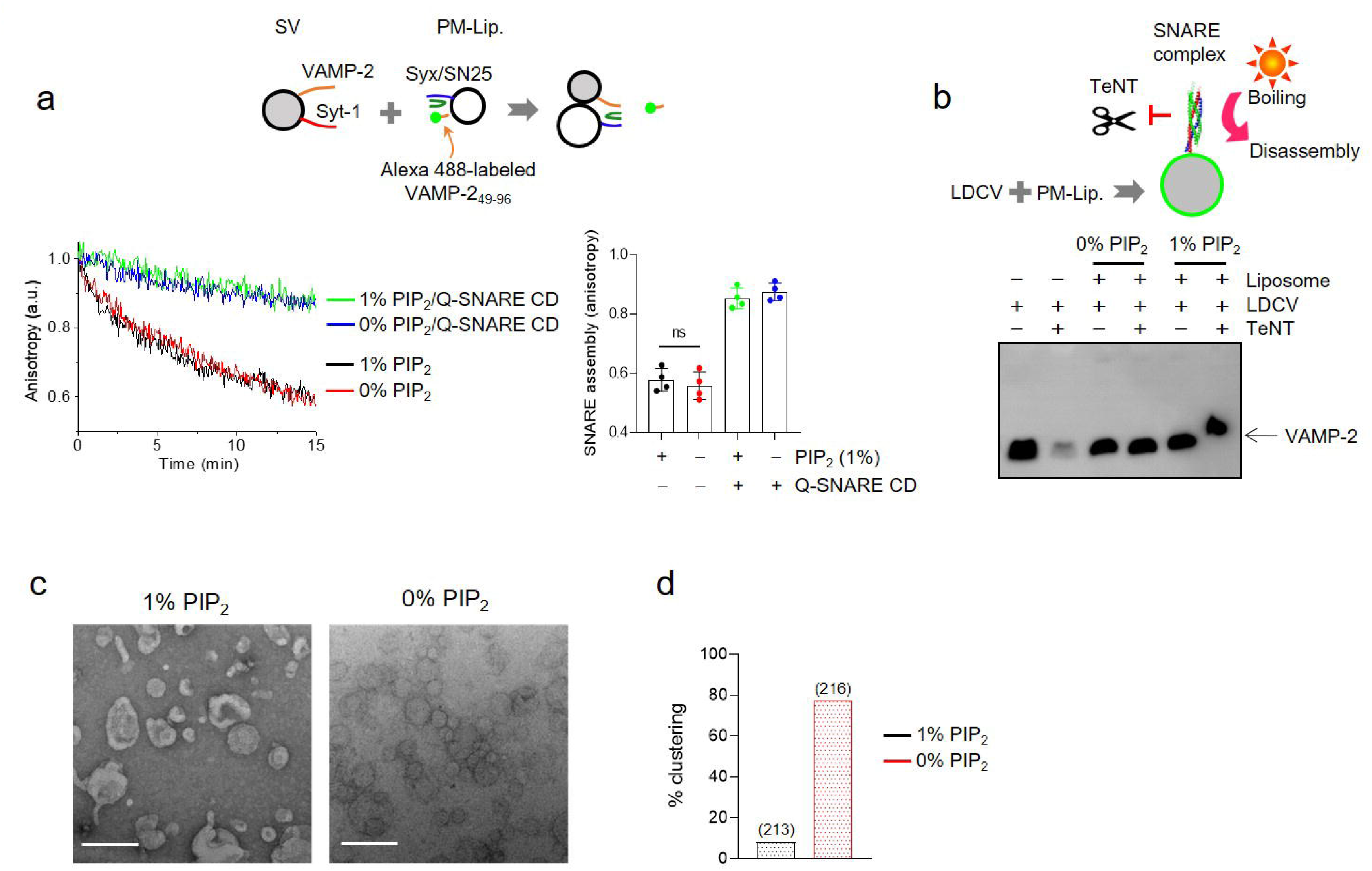
SNARE assembly and vesicle docking in the absence of PIP_2_. (**a**) SNARE assembly was monitored using fluorescence anisotropy. Displacement of the stabilizing VAMP-2 peptide (VAMP-2_49-96_) by endogenous VAMP-2 of native SVs represents SNARE assembly. The stabilized Q-SNARE complex containing VAMP-2_49-96_ labeled with Alexa Fluor 488 (Alexa 488-labeled VAMP-2_49-96_) was incorporated in PM-liposomes. SVs were incubated with PM-liposomes as described in a lipid-mixing assay in Fig. 1b. Preincubation of the Q-SNARE cytoplasmic domain (CD) consisting of syntaxin-1A (aa 183-262) and SNAP-25A with SVs blocked SNARE assembly, and thus prevented dissociation of Alexa 488-labeled VAMP-2_49-96_. Lipid composition of PM-liposomes in **a**: 45% PC, 15% PE, 10% PS, 25% Chol, 4% PI, and 1% PIP_2_. Anisotropy was normalized as A/A_0_, where A_0_ is the initial value of anisotropy. (**b**) Formation of the ternary SNARE complex and SNARE assembly after vesicle fusion either 0% or 1% PIP_2_ in PM-liposomes. LDCVs were incubated with PM-liposomes for 20 min without Ca^2+^ as in Fig. 1a, and then treated with Tetanus neurotoxin (TeNT). TeNT-resistant VAMP-2 indicates the ternary SNARE complex formation in SDS-PAGE. Boiling at 95°C causes the ternary SNARE complex to disassemble, allowing VAMP-2 to migrate to its size. Lipid composition of PM-liposomes is described in Fig. 1. Data are means ± SD from four independent experiments. One-way ANOVA test with Bonferroni correction was used. (**c**) TEM images after LDCV fusion with either 0% or 1% PIP_2_ in PM-liposomes as in Fig. 1a. Scale, 200 nm. (**d**) Quantification of vesicle clustering is presented as a percentage of total vesicles (n = 213, 216 from three independent experiments).

An alternative way to monitor SNARE assembly is to test VAMP-2, which becomes resistant to the light chain of tetanus toxin (TeNT), a protease specific for VAMP-2^46^. Free VAMP-2 of native vesicles is degraded by TeNT, but VAMP-2 engaged in the assembled ternary SNARE complexes is resistant to the cleavage by TeNT; therefore TeNT-resistant VAMP-2 indicates the ternary SNARE complex formation in SDS-PAGE^17, 18, 20, 42^. After LDCV fusion with PM-liposomes as a fusion assay (**Fig. 1a**), TeNT was added to cleave free VAMP-2, and VAMP-2 engaged in the SNARE complex formation was immunoblotted (**Fig. 2b**). As correlating with SNARE assembly using anisotropy (**Fig. 2a**), the ternary SNARE complex formation was observed regardless of PIP_2_ in PM-liposomes (**Fig. 2b**). Altogether, SNAREs are assembled despite no basal fusion of native vesicles in the absence of PIP_2_ in PM-liposomes (**Fig. 1**), suggesting that SNARE complexes are partially zippered in a state of vesicle docking.

To further confirm vesicle docking we carried out negative-stain transmission electron microscopy (TEM) after LDCV fusion with PM-liposomes containing either 0% or 1% PIP_2_ as in **Fig. 1a**. The frequency of vesicle clustering increased when PIP_2_ was excluded from PM-liposomes (**Fig. 2c,d**), showing that fusion is arrested in a state of vesicle docking (0% PIP_2_). These data support that vesicle fusion is arrested at a step after the establishment of the SNARE-mediated docking.

### MARCKS inhibits basal fusion by masking PIP_2_

The experiments described above show that when PM-liposomes have no PIP_2_, Ca^2+^-independent basal fusion of native vesicles is arrested, but partial SNARE zippering leads to vesicle docking. Concentrations of PIP_2_ in a typical mammalian cell ranges from 0.1% to 5% of the inner plasma membrane bilayer lipids^47–50^. Although PIP_2_ is a minor component, it plays a critical role in diverse cellular signaling processes^13^, revealing that the availability of free PIP_2_ is tightly regulated. Myristoylated alanine-rich C-kinase substrate (MARCKS) sequesters and masks PIP_2_ by the electrostatic interaction, and dissociation of MARCKS by Ca^2+^/CaM leads to unmasking of PIP_2_^24, 29^. Given that decreasing PIP_2_ concentrations in PM-liposomes reduces basal fusion of native vesicles (**Fig. 1c**), we investigated whether MARCKS inhibits vesicle fusion by masking PIP_2_.

First of all, MARCKS bound to anionic phospholipids to mask PIP_2_ (**Fig. 3a**). The effector domain of MARCKS (ED, aa 151-175, KKKKKRFSFKKSFKLSGFSFKKNKK) is a fragment of 25 amino acids and enriched with 13 basic residues responsible for the electrostatic interaction with PIP_2_ and 5 aromatic hydrophobic residues for membrane insertion^29^. MARCKS ED binding to PIP_2_ was monitored using FRET measurement in which MARCKS ED was labeled with BODIPY TR at the N-terminus as an acceptor, and Oregon Green-PE was incorporated in protein-free liposomes as a donor dye. Membrane binding of BODIPY TR-labeled MARCKS ED results in quenching of donor fluorescence in liposomes. Anionic phospholipids such as PS and PIP_2_ induced membrane binding of ED electrostatically, because membranes containing no PS/PIP_2_ failed to interact with ED (**Fig. 3a**). Anisotropy measurement confirmed that 1% PIP_2_ alone was enough to cause membrane binding of ED (**Supplementary Fig. 2a**), and thereby confirmed that MARCKS ED electrostatically masks PIP_2_.

**Figure 3.**
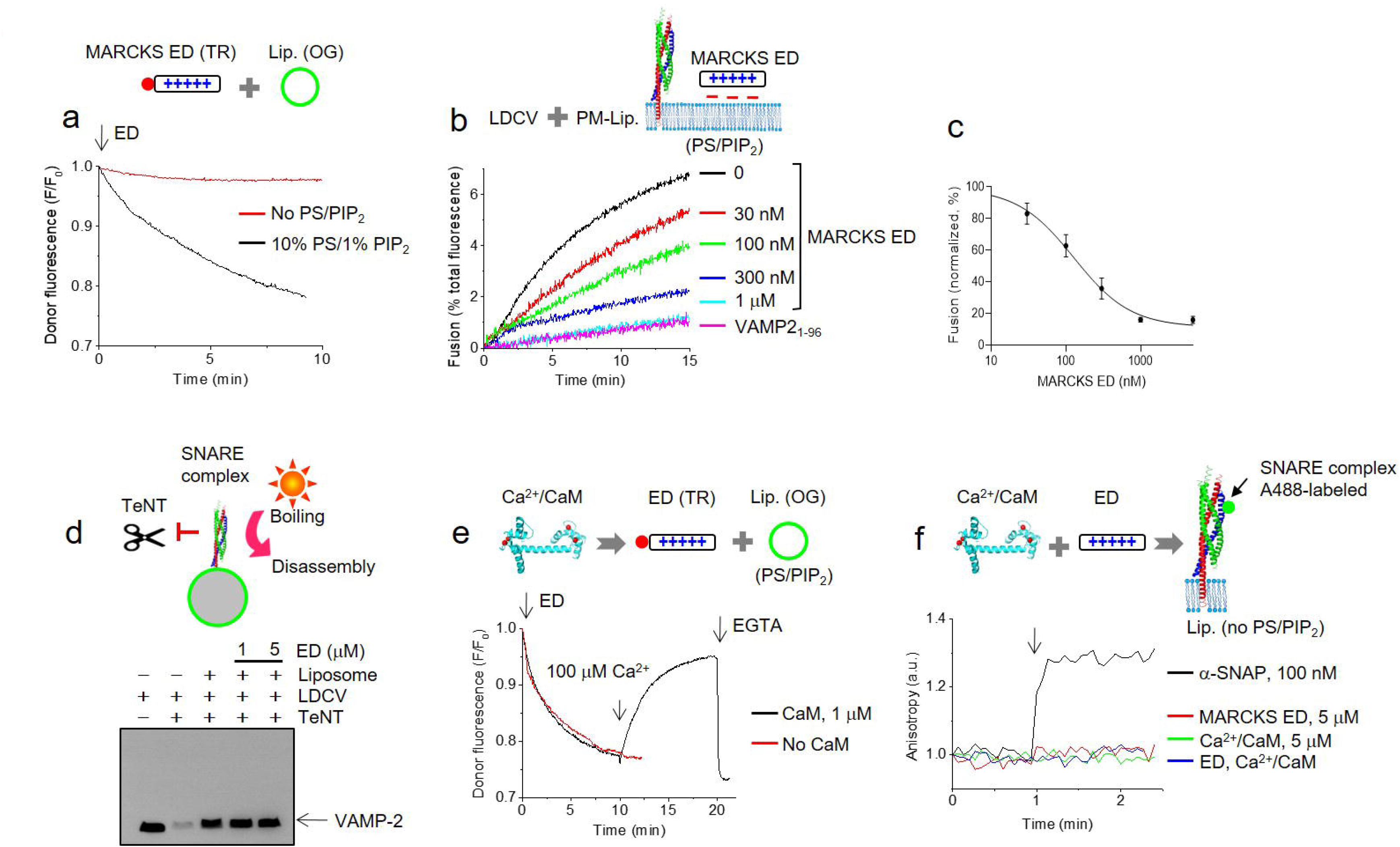
Masking PIP_2_ by MARCKS inhibits vesicle fusion. (**a**) Membrane binding of MARCKS was monitored using FRET measurement. The effector domain (ED) of MARCKS (aa 151–175) was labelled with BODIPY TR and liposomes (lip., protein-free) incorporated Oregon Green (OG) as an acceptor and a donor dye, respectively. Lipid composition of liposomes: 45% PC, 13.5% PE, 1.5% OG 488-DHPE, 10% PS, 25% Chol, 4% PI, and 1% PIP_2_. When PS and PIP_2_ were removed (no PS/PIP_2_), PC contents were adjusted accordingly. 1 mM EGTA was included. (**b,c**) LDCV fusion with PM-liposomes as in Fig. 1a. MARCKS ED (aa 151–175) blocked basal fusion of LDCVs in a dose-dependent manner. Data are means ± SD from three to five independent experiments. IC_50_, 127.6 nM; Hill-slope, 1.143. (**d**) Even in the presence of MARCKS ED, which fully interfered with LDCV basal fusion, SNARE assembly and ternary SNARE complex formation occurred. As in Fig. 2b, TeNT-resistant VAMP-2 represents the ternary SNARE complex formation after a LDCV fusion assay. (**e**) FRET was conducted to monitor association and disassociation of MARCKS ED with liposomes as in Fig. 3a. 100 µM free Ca^2+^ with 1 µM CaM dissociated MARCKS ED from liposomes containing PS/PIP_2_. (**f**) Interaction of the SNARE complex with MARCKS ED and Ca^2+^/CaM was monitored using anisotropy measurement. As in Fig. 2a, the stabilized Q-SNARE complex containing VAMP-2_49-96_ labeled with Alexa Fluor 488 (Alexa 488-labeled VAMP-2_49-96_) was incorporated in liposomes that contained no PS/PIP_2_. Lipid composition of liposomes in **f**: 60% PC, 15% PE, and 25% Chol. Physiological ionic strength with 1 mM MgCl_2_/3 mM ATP was used in all experiments.

MARCKS ED inhibited Ca^2+^-independent LDCV fusion in a dose-dependent manner (**Fig. 3b,c**); IC_50_ = 127.6 nM, Hill-Slope = 1.143. 1 µM ED completely blocked basal fusion, reminiscent of no basal fusion without PIP_2_ in PM-liposomes (**Fig. 1a**).

SNARE assembly and the ternary SNARE complex formation were observable despite no basal LDCV fusion in the presence of 1 µM or 5 µM MARCKS ED (**Fig. 3d**); this result correlates with SNARE assembly in the absence of PIP_2_ in PM-liposomes (**Fig. 2b**).

Ca^2+^/CaM reversed membrane binding of MARCKS ED and dissociated ED from liposomes (**Fig. 3e**). Dynamic and reversible membrane binding of ED was monitored using FRET (**Fig. 3a**). Application of 100 µM Ca^2+^/1 µM CaM dissociated MARCKS ED from PIP_2_-containing membranes, as reported previously^21, 27^. CaM alone without Ca^2+^ bound to ED (**Supplementary Fig. 2b**), but Ca^2+^ was essential for dissociation of ED from PIP_2_-containing membrane, thus Ca^2+^/CaM unmasks PIP_2_ (**Fig. 3d**).

The SNARE complex did not interact with MARCKS ED, Ca^2+^/CaM, or both (**Fig. 3f**), confirming that MARCKS and Ca^2+^/CaM regulate vesicle fusion by masking and unmasking PIP_2_, not by interacting with SNAREs.

### MARCKS prevents membrane binding of the C2AB domain by masking PIP_2_

We have previously reported that PIP_2_ is also required for Ca^2+^-triggered fusion of native vesicles by inducing the *trans*-interaction of synaptotagmin-1 with the plasma membrane; without PIP_2_ in PM-liposomes, Ca^2+^-dependent fusion of native vesicles does not occur^17–19^. Therefore, we investigated whether PIP_2_-masking MARCKS ED reduces synaptotagmin-1 binding to PIP_2_-containing membranes. The C2AB domain (Syt_97-421_) of synaptotagmin-1 was labelled with Alexa Fluor 488 at S342C as a donor, and liposomes (Lip., protein-free, 10% PS/1% PIP_2_ included as in PM-liposomes) were labelled with Rhodamine (Rho)-PE as an acceptor (**Fig. 4**). As a control, we confirmed no interaction of the C2AB domain with MARCKS ED, Ca^2+^/CaM, or both in the absence of liposomes (**Fig. 4a**).

**Figure 4.**
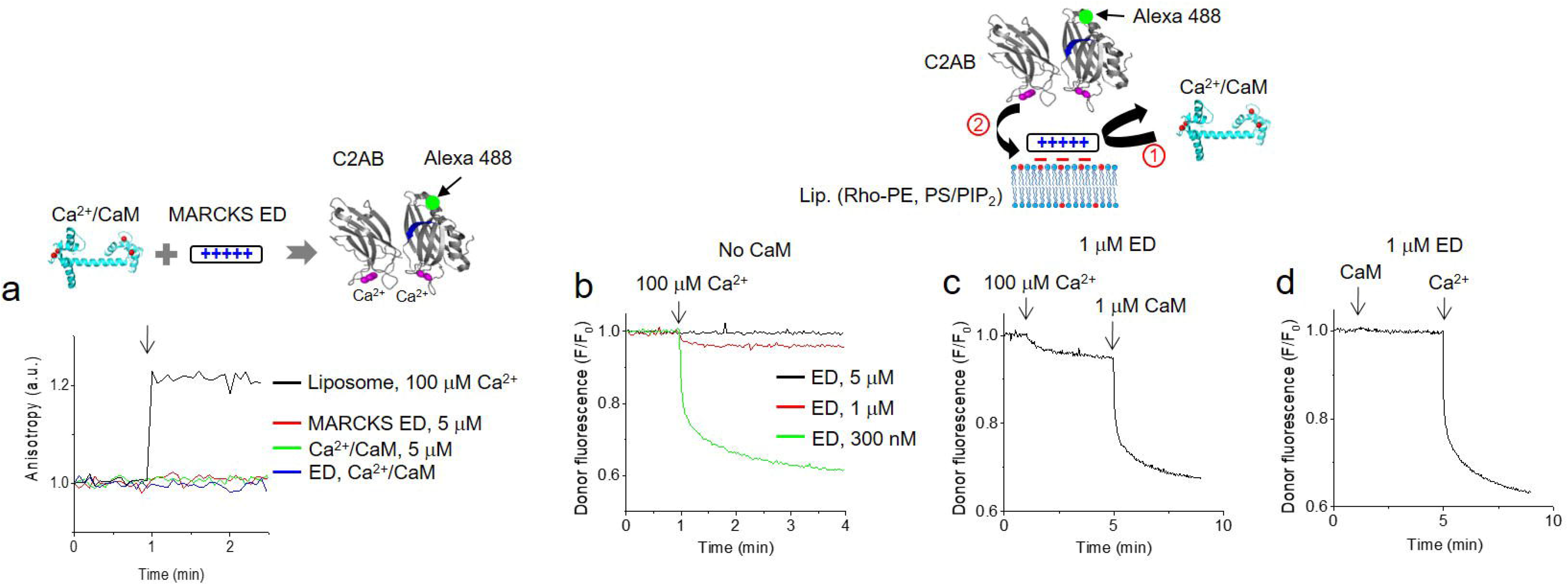
Unmasking PIP_2_ by Ca^2+^/CaM leads to synaptotagmin-1 membrane binding. (**a**) Anisotropy was carried out to monitor the interaction of the C2AB domain of synaptotagmin-1 with MARCKS ED and Ca^2+^/CaM. The C2AB domain (Syt-1_97-421_) was labeled with Alexa Fluor 488 at S342C (green dots). No liposomes were included. (**b-d**) Monitoring the membrane binding of the C2AB domain using FRET measurement. The C2AB domain (Syt-1_97-421_) was labeled with Alexa Fluor 488 at S342C (green dots) as a donor dye, and liposomes (Lip.) incorporated Rhodamine (Rho)-PE (red) as an acceptor dye. Lipid composition of liposomes for FRET: protein-free, 45% PC, 13.5% PE, 1.5% Rho-PE, 10% PS, 25% Chol, 4% PI, and 1% PIP_2_. (**b**) MARCKS ED interfered in a dose-dependent manner with the C2AB membrane binding evoked by 100 µM free Ca^2+^. 1 µM MARCKS ED completely blocked membrane binding of the C2AB domain by 100 µM free Ca^2+^. (**c,d**) In the presence of 1 µM MARCKS ED, 100 µM free Ca^2+^ and 1 µM CaM were sequentially applied to induce the C2AB membrane binding. FRET was normalized as F/F_0_, where F_0_ is the initial value of the donor fluorescence intensity. Physiological ionic strength with 1 mM MgCl_2_/3 mM ATP was used in all experiments.

C2AB binding to liposomes was monitored using FRET between the C2AB domain (Alexa Fluor 488) and Rhodamine-labeled liposomes (**Fig. 4b-d**). In the absence of ED, addition of 100 µM free Ca^2+^ caused robust membrane binding of the C2AB domain, but Ca^2+^/CaM did not affect the C2AB binding to liposomes that contained PIP_2_ (**Supplementary Fig. 3**), because Ca^2+^/CaM has no interaction with the C2AB domain (**Fig. 4a**). As expected, MARCKS ED prevented Ca^2+^-induced membrane binding of the C2AB domain in a dose-dependent manner by masking PIP_2_ (**Fig. 4b**).

In the presence of 1 µM ED, Ca^2+^ and CaM were applied sequentially to rescue the C2AB membrane binding by dissociating ED and unmasking PIP_2_ (**Fig. 4c,d**). Addition of 1 µM ED, which masks PIP_2_, dramatically inhibited membrane binding of the C2AB domain that is induced by 100 µM free Ca^2+^ (**Fig. 4c**). Addition of 100 µM Ca^2+^ alone slightly induced C2AB binding, then sequential treatment of Ca^2+^/CaM accelerated and rescued the C2AB binding (**Fig. 4c**). CaM alone did not rescue the C2AB membrane binding, but Ca^2+^/CaM caused membrane dissociation of ED (**Fig. 3e**), thereby rescuing the C2AB membrane binding by unmasking PIP_2_ (**Fig. 4d)**.

### CaM potentiates Ca^2+^-dependent vesicle fusion mediated by synaptotagmin-1

The data presented above suggest that MARCKS ED inhibits Ca^2+^-dependent membrane binding of the C2AB domain by masking PIP_2_, and Ca^2+^/CaM recovers the C2AB membrane binding by dissociating ED and unmasking PIP_2_ (**Fig. 3e**). Therefore, we tested whether Ca^2+^/CaM accordingly increases Ca^2+^-dependent vesicle fusion by unmasking PIP_2_. Addition of 1 µM ED completely inhibited LDCV basal fusion to the level of VAMP-2_1-96_ treatment (**Fig. 3b**), and 1 µM CaM without Ca^2+^ in the presence of ED failed to rescue basal fusion (**Fig. 5a**), because CaM without Ca^2+^ had no effect on membrane dissociation of ED (**Fig. 3e**). In the presence of 1 µM ED, but the absence of CaM, 100 µM Ca^2+^ alone slightly induced Ca^2+^-dependent vesicle fusion (**Fig. 5a**), correlating with the weak C2AB binding to liposomes (100 µM Ca^2+^, 1 µM ED, **Fig. 4b,c**). Indeed, Ca^2+^/CaM potentiated Ca^2+^-dependent vesicle fusion (**Fig. 5a**); this result correlates with the ability of Ca^2+^/CaM to rescue the C2AB membrane binding (**Fig. 4c,d**).

**Figure 5.**
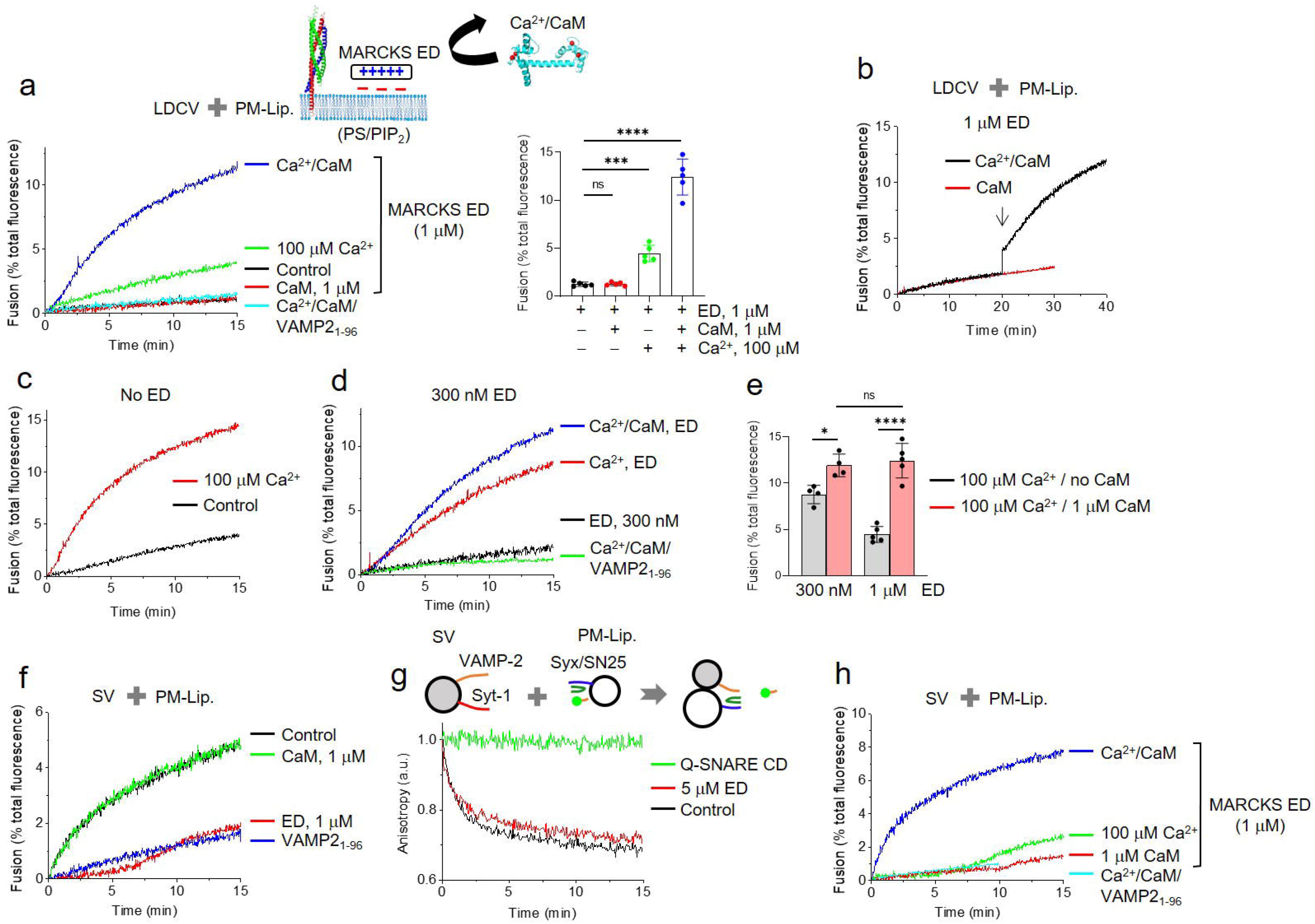
Ca^2+^/CaM triggers vesicle fusion arrested by MARCKS ED. (**a-e**) LDCV fusion with PM-liposomes using a lipid-mixing assay as in Fig. 1a. VAMP-2_1-96_, the cytoplasmic domain of VAMP-2 (aa 1-96), fully disrupted SNARE-dependent fusion. 1 µM MARCKS ED completely inhibited LDCV basal fusion. 100 µM free Ca^2+^ alone slightly recovered LDCV fusion, but CaM potentiated Ca^2+^-dependent LDCV fusion. (**b**) Sequential treatment of ED and Ca^2+^/CaM during fusion reaction. (**c**) No MARCKS ED was included. 100 µM Ca^2+^ accelerated LDCV fusion. (**d,e**) In the presence of 300 nM MARCKS ED, Ca^2+^ alone accelerated LDCV fusion, and CaM potentiated Ca^2+^-dependent LDCV fusion. Data are means ± SD from four or five independent experiments. One-way ANOVA test with Bonferroni correction was used; *, *p* < 0.05, ****, *p* < 0.0001. (**f**) SV fusion with PM-liposomes (Fig. 1b). (**g**) Monitoring SNARE assembly using fluorescence anisotropy as in Fig. 2a. Endogenous VAMP-2 of native SVs displaces Alexa 488-labeled VAMP-2_49-96_ by SNARE assembly when SVs fuse with PM-liposomes. Lipid composition in **g**: 45% PC, 15% PE, 10% PS, 25% Chol, 4% PI, and 1% PIP_2_. Anisotropy was normalized as A/A_0_, where A_0_ is the initial value of anisotropy. (**h**) SV fusion with PM-liposomes in the presence of 1 µM MARCKS ED. Ca^2+^ alone slightly triggered SV fusion, and CaM potentiated Ca^2+^-dependent SV fusion. Physiological ionic strength with 1 mM MgCl_2_/3 mM ATP was used in all experiments.

Next, we tested whether Ca^2+^/CaM can trigger LDCV fusion, which is arrested by ED in a state of vesicle docking through the partially zippered SNARE complex (**Fig. 2**). Addition of 1 µM ED was used to arrest LDCV basal fusion by masking PIP_2_, then Ca^2+^/CaM was applied in a state of docking of LDCVs with PM-liposomes; Ca^2+^/CaM triggered LDCV fusion like a step increase, but CaM alone was ineffective (**Fig. 5b**). Taken together, these results indicate that Ca^2+^/CaM unmasks PIP_2_ by dissociating MARCKS ED, and thereby potentiates Ca^2+^-dependent vesicle fusion, because the C2AB domain of synaptotagmin-1 requires PIP_2_ to trigger fusion^17, 18, 20^.

PIP_2_ actively interacts with partner proteins for various cellular functions^13^, so free PIP_2_ is tightly regulated in cells. Therefore, we tested MARCKS ED at low concentrations, at which free PIP_2_ is available for basal fusion. In the absence of ED, LDCVs underwent robust basal fusion, and 100 µM Ca^2+^ accelerated LDCV fusion (**Fig. 5c**). Addition of 1 µM ED completely arrested basal LDCV fusion (**Fig. 3b,5a**). In the presence of 300 nM ED, basal fusion was slightly rescued (**Fig. 3b,5d**); addition of only Ca^2+^ accelerated LDCV fusion, and Ca^2+^/CaM potentiated Ca^2+^-dependent LDCV fusion (**Fig. 5d,e**). Increasing ED concentrations strongly inhibited Ca^2+^-dependent fusion, but Ca^2+^/CaM rescued it (**Fig. 5a,d,e**).

### SVs reproduce Ca^2+^-dependent vesicle fusion potentiated by CaM

We further tested SV fusion to obtain generalized conclusions that Ca^2+^/CaM potentiates Ca^2+^-dependent fusion. Addition of 1 µM MARCKS ED completely inhibited SV basal fusion to the level of VAMP-2_1-96_ treatment by PIP_2_-masking (**Fig. 1b, Fig. 5f**); this is similar to the result obtained using LDCVs (**Fig. 5a**). CaM alone had little effect on basal SV fusion (**Fig. 5f**).

We also monitored SNARE assembly by using fluorescence anisotropy (**Fig. 2a**) to confirm whether SV fusion is arrested in a state of docking, in which SNARE proteins are assembled. As expected, even in the presence of 5 µM ED that fully arrested basal SV fusion (**Fig. 5f**), SNARE zippering and assembly occurred (**Fig. 5g**); this result is similar to SNARE assembly of LDCVs independent of PIP_2_ (**Fig. 2b,3d**).

Addition of 100 µM Ca^2+^ in the presence of 1 µM ED slightly induced Ca^2+^-dependent vesicle fusion, but Ca^2+^/CaM potentiated Ca^2+^-dependent SV fusion by dissociating MARCKS and unmasking PIP_2_ (**Fig. 5h**); this process is consistent with LDCV fusion. As a control, we tested whether CaM might synergize with synaptotagmin-1 to induce Ca^2+^-dependent vesicle fusion independent of MARCKS. In the absence of MARCKS ED, 100 µM Ca^2+^ accelerated SV fusion, but Ca^2+^/CaM had no effect on Ca^2+^-dependent SV fusion mediated by synaptotagmin-1 (**Supplementary Fig. 4**). It occurs because Ca^2+^/CaM has no interaction with SNAREs (**Fig. 3f**), the C2AB domain (**Fig. 4a**), or membranes (**Supplementary Fig. 3**), indicating that CaM potentiates Ca^2+^-dependent SV fusion by unmasking PIP_2_. Altogether, free PIP_2_, which is dynamically regulated by MARCKS and Ca^2+^/CaM, is the key to trigger both basal fusion and Ca^2+^-dependent fusion of native vesicles.

### Unmasking of PIP_2_ by PKC-epsilon causes Ca^2+^-independent basal fusion

MARCKS electrostatically masks PIP_2_, and Ca^2+^/CaM unmasks PIP_2_ by dissociating MARCKS from PIP_2_-containing membranes (**Fig. 3a,e**), thus leading to potentiation of Ca^2+^-dependent vesicle fusion (**Fig. 5**). CaM dissociates MARCKS from PIP_2_ in a Ca^2+^-dependent manner (**Fig. 3e**). In addition to Ca^2+^-dependent fusion, free PIP_2_ is required for Ca^2+^-independent basal fusion of native vesicles (**Fig. 1, Supplementary Fig. 1**). Basal fusion and some of spontaneous release *in-vivo* are Ca^2+^-independent, so we hypothesize that Ca^2+^-independent membrane dissociation of MARCKS ED can trigger spontaneous release and basal fusion.

PKC dissociates MARCKS from PIP_2_ by phosphorylation^25, 29^. PKC-epsilon, an isoform of novel PKC subfamily, is activated by diacylglycerol (DAG), but not by Ca^2+51^. In the final set of experiments, we tested whether membrane dissociation of MARCKS ED by PKC-epsilon induces basal fusion of LDCVs. We monitored membrane dissociation of ED by using FRET (**Fig. 3a,e**). PKC-epsilon dissociated ED only in the presence of phorbol 12-myristate 13-acetate (PMA), an analog of DAG and activator of PKC (**Fig. 6a,b**). Activation of PKC-epsilon resulted in triggering of basal LDCV fusion, which was completely arrested by 1 µM ED (**Fig. 6c**).

**Figure 6.**
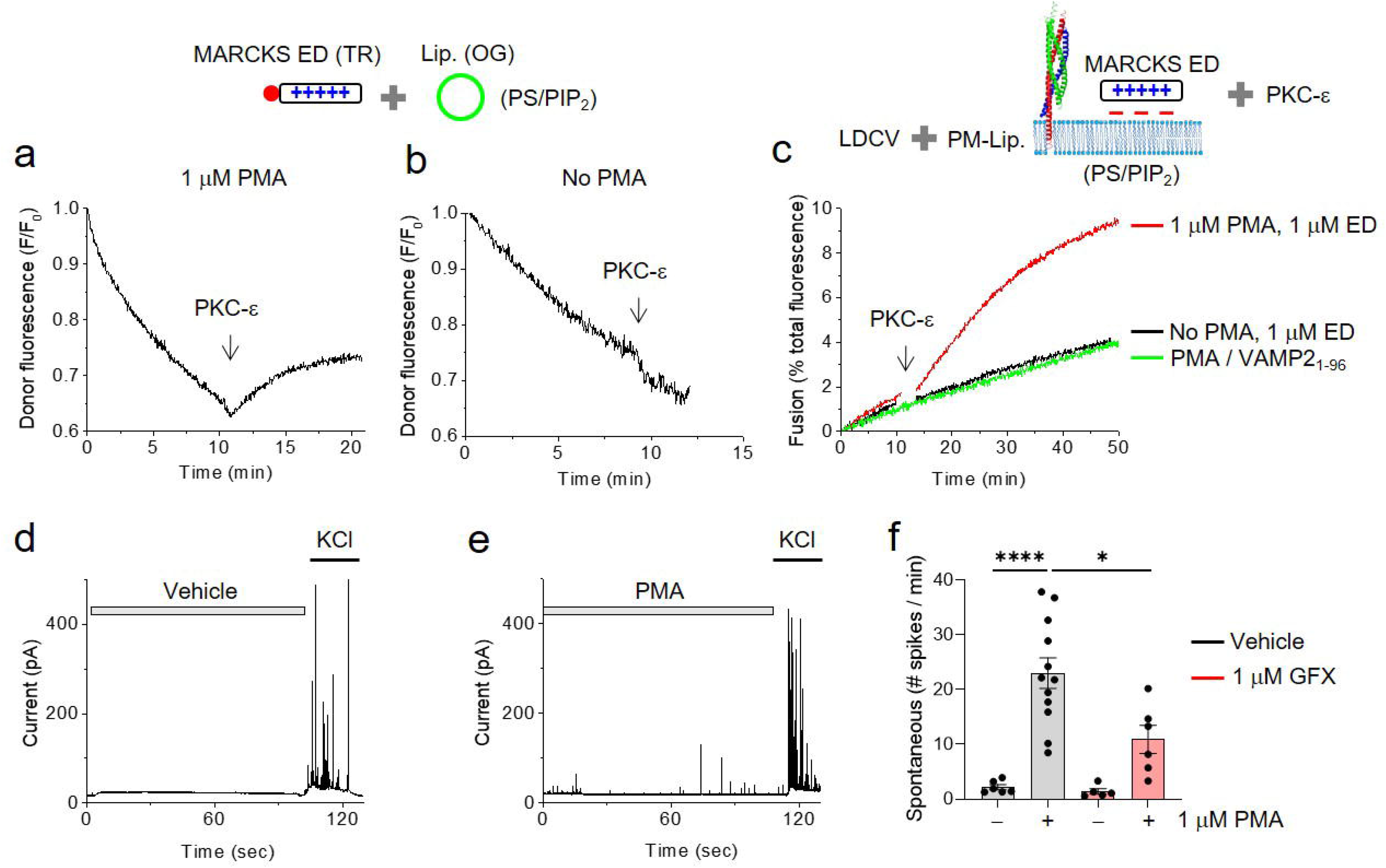
PKC-epsilon dissociates MARCKS ED and rescues vesicle fusion. (**a,b**) Monitoring membrane association and disassociation of MARCKS ED using FRET measurement as in Fig. 3a. 60 nM PKC-epsilon was applied in the presence (**a**) or absence (**b**) of 1 µM PMA. 10% PS/1% PIP_2_ were included in lip. (**c**) LDCV fusion using a lipid-mixing assay as in Fig. 1a. PKC-epsilon with PMA recovered LDCV fusion. Physiological ionic strength with 1 mM MgCl_2_/3 mM ATP was used in all experiments; ATP removes contaminating Ca^2+19^. (**d-f**) Monitoring LDCV exocytosis in bovine chromaffin cells using amperometry. Shown are typical amperometric traces of chromaffin cells pre-treated for 5 min with DMSO vehicle (**d**) or 1 µM PMA (**e**). 50 mM KCl stimulations were applied to confirm functional exocytosis of chromaffin cells. (**f**) Spontaneous LDCV exocytosis before KCl stimulation is presented as numbers of amperometric spikes per min. Data are means ± SEM from three independent experiments (n = 6, 12, 5, 6). One-way ANOVA test with Bonferroni correction was used; *, *p* < 0.05, ****, *p* < 0.0001.

PKC activation increased spontaneous release of LDCVs in chromaffin cells. To test the PKC effect on basal fusion, we used amperometry to monitor spontaneous LDCV exocytosis in real time. Treatment with PMA increased spontaneous release of LDCVs, which was reduced by GF109203X (GFX), an inhibitor of PKC (**Fig. d-f**). Taken together, these results suggest that ED dissociation by PKC-epsilon triggers Ca^2+^-independent basal fusion by unmasking PIP_2_.

### Mathematical modeling to explain how PIP_2_ triggers vesicle fusion

To investigate the molecular mechanisms by which PIP_2_ electrostatically triggers Ca^2+^-independent vesicle fusion, we used mathematical modeling with the modified Derjaguin-Landau-Verwey-Overbeek (DLVO) theory suggested by Ohki and Oshima^52^ (**Supplementary Fig. 5,6**). PIP_2_ might lower the hydration energy, because PIP_2_ results in accumulation of cations such as K^+^ and Mg^2+^, which brings two anionic membranes more closely by neutralizing electrostatic repulsion^53–55^. During vesicle docking, the van der Waals attraction and hydrophobic interactions from hydrophobic lipid tails increase^53^ (**Supplementary Fig. 6**), and thereby promote vesicle fusion.

## Discussion

We have previously reported that α-<u>s</u>oluble <u>N</u>SF <u>a</u>ttachment <u>p</u>rotein (α-SNAP), the dominant isoform of SNAPs, binds to the Q-SNARE complexes and interferes with full SNARE zippering, thus slowing down fusion and causing tight docking as a result of partially assembled SNARE proteins^42^. The SNARE complex is disassembled by the AAA+ ATPase *N*-ethylmaleimide-sensitive factor (NSF) and a cofactor, α-SNAP. α-SNAP first binds to the SNARE complex with a high binding affinity, and then recruits NSF, thereby catalyzing SNARE disassembly in an ATP-dependent manner^56^. α-SNAP that binds to the Q-SNARE complexes induces partial SNARE zippering in a state of tight docking of LDCVs^42^. Importantly, α-SNAP dramatically inhibits LDCV (∼150 nm in diameter) fusion within 5 min after fusion reaction, but does not significantly affect SV (∼45 nm in diameter) fusion, because high curvature of small SVs is likely to overcome the inhibitory effect of α-SNAP^42^. The curvature of vesicle membrane affects the energy landscape during fusion reaction, so arresting SNARE assembly of small SVs is energetically challenging^42^. The arrest and clamping of SNARE assembly during vesicle docking and priming has been hypothesized, although the proteins proposed to cause this arrest, such as complexin and synaptotagmin-1, remain controversial and unclear^4^.

Here we provide evidence that full SNARE zippering and basal fusion of both LDCVs and SVs are arrested in a state of vesicle docking. PIP_2_ masking by MARCKS causes vesicle fusion to be arrested and halted, even though SNAREs are partially assembled (**Fig. 2, 3**). Ca^2+^/CaM dissociates MARCKS, then unmasks PIP_2_, thereby potentiating synaptotagmin-1-mediated Ca^2+^-dependent vesicle fusion (**Fig. 4, 5**). MARCKS can also be dissociated to unmask PIP_2_ independently of Ca^2+^ by activating PKC-epsilon, leading to Ca^2+^-independent basal fusion (**Fig. 6**). Masking PIP_2_ results in partial zippering of SNAREs and arresting fusion of both LDCVs and SVs, indicating that free PIP_2_ has a critical function in both Ca^2+^-dependent and Ca^2+^-independent vesicle fusion. This mechanism is unlike α-SNAP, which interferes only with SNARE zippering of LDCVs^42^.

Purified native vesicles, LDCVs and SVs, have significant advantages for the *in-vitro* reconstitution of vesicle fusion in a physiological context. V-liposomes are not dependent on PIP_2_ for fusion, and fail to reproduce arrest of vesicle fusion by masking PIP_2_; this observation suggests that the reconstitution of physiological vesicle fusion requires purified native vesicles. V-liposomes may have limitations to replace native vesicles; however, it is currently unclear what distinguishes native vesicles from V-liposomes. The structural integrity of native vesicles, such as lipid composition, protein density, and membrane physical property, allows them to mimic endogenous vesicle fusion in the *in-vitro* reconstitution. This observation is similar to a previous report on the cholesterol effect^20^. Compared to V-liposomes, native vesicles have higher membrane rigidity and seem to require more energy to overcome the barrier for fusion^20^. Native vesicle fusion is arrested and halted without PIP_2_, and native vesicles require PIP_2_-induced electrostatic attraction to overcome this energy barrier (**Fig. 7a**). However, V-liposomes may have a low energy barrier for fusion, so PIP_2_-induced electrostatic attraction may not be necessary for overcoming this energy barrier. Increasing the amount of PIP_2_ facilitates liposome-liposome fusion, suggesting PIP_2_ increases membrane fusion by lowering the energy barrier (**Fig. 1d**).

**Figure 7.**
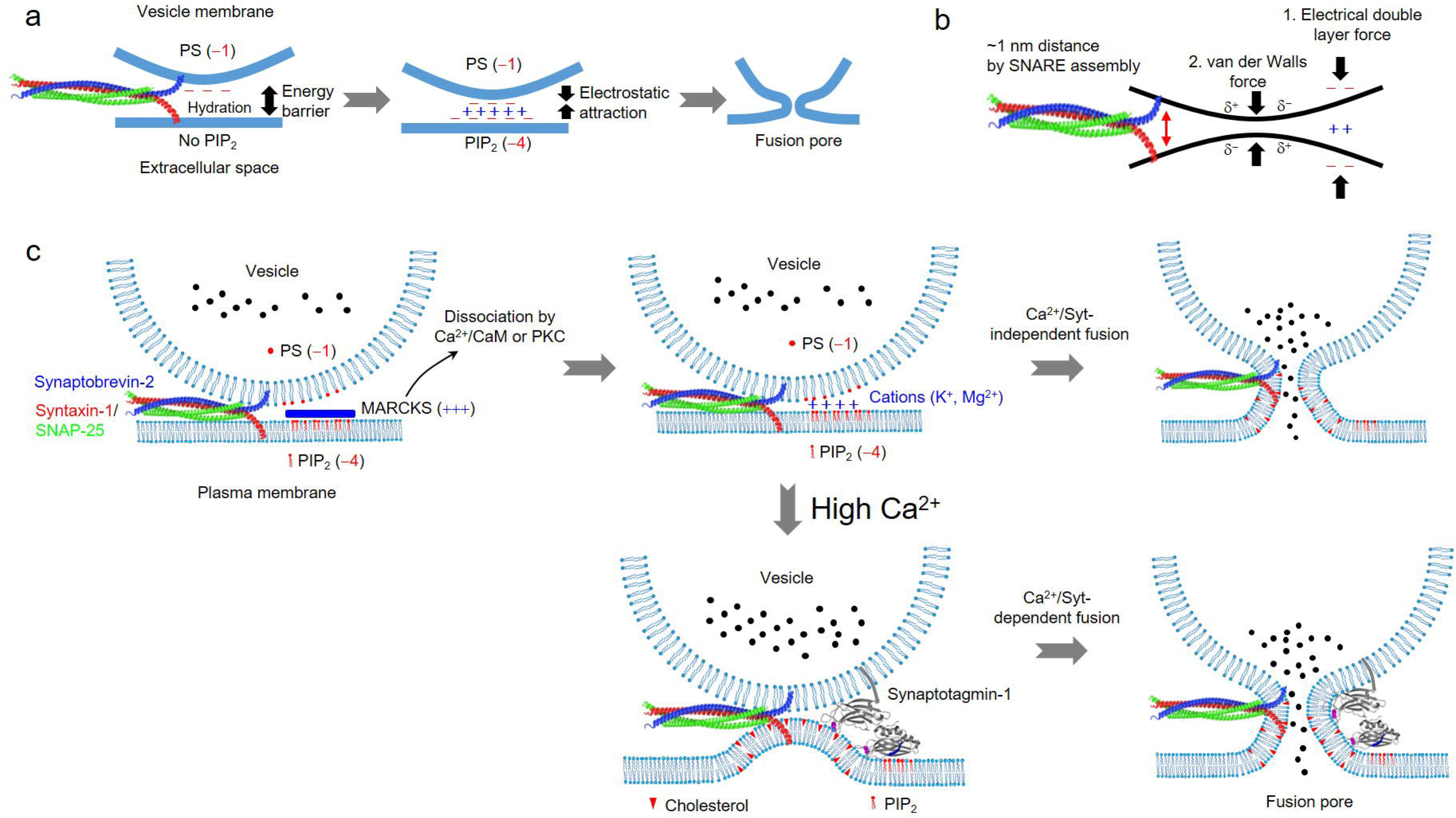
Schematic diagram of proposed new paradigm in which free PIP_2_ is a triggering factor of vesicle fusion by generating electrical double layer forces. (**a**) Free PIP_2_ electrostatically triggers vesicle fusion by overcoming an energy barrier. PS (−1 net charge at pH 7) is present in vesicle membranes and PIP_2_ (−4 net charge at pH 7) is present in the plasma membranes; MARCKS masks PIP_2_. When two membranes approach ∼1 nm distance by the partial SNARE assembly, the energy barrier might arrest vesicle fusion. Unmasking PIP_2_ causes cations to accumulate, including K^+^ and Mg^2+^, which bring these two anionic membranes into close proximity by neutralizing electrostatic repulsion, potentially leading to vesicle fusion. (**b**) According to the Derjaguin-Landau-Verwey-Overbeek (DLVO) theory, (1) electrical double layer forces attract two membranes very closely on the order of a few Ångstroms. (2) the van der Waals attraction increases as two membranes get closer, and can overcome the energy barrier for two membranes to fuse. (**c**) Free PIP_2_ is required for both Ca^2+^-dependent and Ca^2+^-independent vesicle fusion. Free PIP_2_ electrostatically triggers Ca^2+^-independent fusion by generating the electrical double layer forces. For Ca^2+^-dependent fusion, PIP_2_ causes the *trans*-interaction of synaptotagmin-1 with the plasma membrane, and synaptotagmin-1 triggers vesicle fusion by membrane bending, which is strengthened by cholesterol.

Our data show that vesicle fusion requires dynamic regulation of free PIP_2_ by CaM and PKC. In glutamatergic neurons, CaM is involved in Ca^2+^-dependent spontaneous release^57^, supporting our model that Ca^2+^/CaM increases basal fusion by unmasking PIP_2_. Phorbol esters increase Ca^2+^-independent spontaneous release through Munc13 and PKC^58^ in inhibitory and excitatory synapses^59–61^, again supporting that PKC-epsilon induces basal fusion in a Ca^2+^-independent manner by unmasking PIP_2_ (**Fig. 6**). In neurons, CaM and PKC are essential signalling proteins, so their interplay potentiates Ca^2+^-dependent exocytosis by modulating membrane-attached proteins^36^. Our data suggest a novel mechanism by which PKC increases spontaneous release, and Ca^2+^/CaM potentiates Ca^2+^-dependent vesicle fusion by unmasking PIP_2_.

Force measurements of SNARE complex unzipping show the presence of an energy barrier in the midpoint of the SNARE complex formation^62, 63^. α-SNAP, which has a high binding affinity for SNAREs, might elevate an energy barrier, and thereby prevent full SNARE zippering. α-SNAP interferes with SNARE zippering and thus slows LDCV fusion, but LDCV fusion is gradually rescued^42^. However, α-SNAP has little effect on SV fusion^42^.

Synaptotagmin-1, which is a candidate for SNARE clamping factors, can interact with SNAREs, but only at low ionic strength, and Mg^2+^/ATP almost completely disrupts this extremely weak synaptotagmin-1−SNAREs interaction by the charge shielding effect as reported previously^18, 64^. Overall, SNARE zippering and vesicle fusion are rarely arrested by SNARE-interacting proteins in a state of tight docking.

How can masking PIP_2_ arrest full SNARE zippering and vesicle fusion? Repulsive energy barriers, for instance the hydration energy barrier, can arrest native vesicle fusion^65^. Water molecules are tightly bound to the surface of the lipid bilayers, and this hydration layer prevents vesicle fusion^65^. The hydration energy barrier must be overcome by a combination of mechanical forces, hydrophobic and electrostatic interactions^65^. Water molecules are displaced from the membrane interface, thus allowing two membranes to come into close contact and fuse together. Interaction of cations such as K^+^ and Mg^2+^ with the polar head groups of phospholipids facilitates displacement of water molecules from the membrane interface^54^.

Vesicle membranes contain PS (−1 net charge at pH 7) and the plasma membranes have PIP_2_ (−4 net charge at pH 7)^12^, which is masked by MARCKS (**Fig. 7a**). When two membranes approach ∼1 nm distance by the partial SNARE assembly^9, 10^, the repulsive hydration force might arrest vesicle fusion by preventing efficient membrane fusion^66^. Unmasking PIP_2_ causes cations to accumulate, including K^+^ and Mg^2+^, which bring these two anionic membranes more closely by neutralizing electrostatic repulsion^54, 55^. Cations can form bridges between the negatively-charged head groups on opposing membranes, thereby promoting their close association and fusion^67^ (**Fig. 7a**). Cations repel water molecules, and thus increase the hydrophobicity, lowering the hydration energy (**Supplementary Fig. 6**). We propose that free PIP_2_ triggers vesicle fusion electrostatically by reducing the hydration energy (**Supplementary Fig. 5,6**).

According to the Derjaguin-Landau-Verwey-Overbeek (DLVO) theory, electrical double layer forces attract two membranes into close proximity when SNAREs are assembled, but electrostatic repulsion begins to increase^53^ (**Fig. 7b**). However, the van der Waals attraction and the hydrophobic interactions from hydrophobic lipid tails also increase as two membranes approach each other^53^ (**Fig. 7b**). The van der Waals and hydrophobic interactions cause two anionic membranes to have an attractive force when the distance between the membranes is on the order of a few Ångstroms; the van der Waals forces become strong enough to overcome the energy barrier, thereby inducing two membranes to fuse^53^. Despite strong experimental evidences, further investigation using molecular dynamics simulation must be conducted to determine how free PIP_2_ electrostatically triggers vesicle fusion.

Free PIP_2_ is required for both Ca^2+^-dependent and Ca^2+^-independent vesicle fusion (**Fig. 7c**). Free PIP_2_ electrostatically triggers Ca^2+^-independent fusion by generating the electrical double layer forces that eventually reduce the hydration energy barrier. For Ca^2+^-dependent fusion, PIP_2_ causes the *trans*-interaction of synaptotagmin-1 with the plasma membrane^17, 18^, and synaptotagmin-1 triggers vesicle fusion by membrane bending, which is strengthened by cholesterol^20^. Ca^2+^ fails to trigger synaptotagmin-1-mediated Ca^2+^-dependent fusion when PIP_2_ is removed from PM-liposomes^17^, and PIP_2_ critically regulates Ca^2+^-sensitivity and Ca^2+^-cooperativity of synaptotagmin-1^19^. Overall, our findings provide novel mechanisms and suggest a new paradigm in which free PIP_2_ is a triggering factor of vesicle fusion at physiological ionic strength by generating the electrical double layer forces.

## Data availability

The datasets that support the findings of this study are openly available.

## Acknowledgements

We thank Dr. Reinhard Jahn for constructs and samples, and to Dr. Kyong-Tai Kim for technical assistanace. Thanks to Dr. Ahmed Elalawy and Sarra Karrar from Widam Food Company for the arrangement of adrenal glands. Thanks to HBKU Core Labs for technical supports. This work was supported by the grant from Qatar Biomedical Research Institute (Project Number SF 2019 004 and IGP5-2022-001 to Y.P.) and the HBKU Thematic Research Grant (Project Number VPR-TG02-06 to Y.P.).

## Author Contributions

Conceptualization, Y.P.; method, investigation, and analysis, Y.P., H.Y.A.M., K.C.S., J.P.; supervision, Y.P., S.M.; project administration and funding acquisition, Y.P.; mathematical modelling, S.H.P., O.S.L.; original draft preparation, Y.P.; review and editing, H.Y.A.M., K.C.S., J.P., S.M., S.H.P., O.S.L.; All authors have read and agreed to the published version of the manuscript.

---

**Supplementary Figure 1 No LDCV fusion in the absence of PIP_2_ in PM-liposomes. (a,b)** A LDCV fusion assay as in Fig. 1a. PM-liposomes contained either 0% PIP_2_ or 1% PIP_2_. PM-liposomes prepared by the standard method (small unilamellar vesicles, SUVs) were incubated with LDCV (**a**) or SV (**b**). (**c**) PM-liposomes incorporated the Q-SNARE complex consisting of the full-length syntaxin-1A (1-288) and SNAP-25A (no cysteine, cysteines replaced by alanines). Lipid composition of PM-liposomes: 45% PC, 12% PE, 3% labeled PE, 10% PS, 25% Chol, 4% PI, and 1% PIP_2_. When PIP_2_ was removed (0% PIP_2_), PI contents were adjusted accordingly. VAMP-2_1-96_, the cytoplasmic domain of VAMP-2 (aa 1-96), blocked SNARE-dependent vesicle fusion. (**d**) The size distribution of proteoliposomes was determined using dynamic light scattering (DLS). SUVs (∼50 nm in diameter) were prepared by the standard method.

**Supplementary Figure 2 Binding of MACRKS ED to PIP_2_-containing membrane and CaM. (a)** Monitoring the membrane binding of MARCKS ED using fluorescence anisotropy. MARCKS ED (aa 151–175) was labeled with BODIPY 493/503. Lipid composition of liposomes (protein-free): 74% PC, 25% Chol, and 1% PIP_2_. When PIP_2_ was removed (no PIP_2_), PC contents were adjusted accordingly. (**b**) Monitoring the interaction of MARCKS ED with CaM using anisotropy. ED binds to CaM and PS/PIP_2_-containing membrane. MARCKS ED was labeled with BODIPY TR. Lipid composition of liposomes (protein-free): 45% PC, 15% PE, 10% PS, 25% Chol, 4% PI, and 1% PIP_2_. When PS and PIP_2_ were removed (no PS/PIP_2_), PC contents were adjusted accordingly. Ca^2+^ was omitted. Physiological ionic strength with 1 mM MgCl_2_/3 mM ATP was used in all experiments.

**Supplementary Figure 3 No effect of CaM on Ca^2+^-dependent C2AB membrane binding. (a)** FRET measurement was applied to monitor the membrane binding of the C2AB domain as described in Fig. 4b. The C2AB domain (Syt-1_97-421_) was labelled with Alexa Fluor 488 at S342C (green dots) as a donor dye, and liposomes (Lip.) incorporated Rhodamine (Rho)-PE (red) as an acceptor dye. Lipid composition of liposomes for FRET: protein-free, 45% PC, 13.5% PE, 1.5% Rho-PE, 10% PS, 25% Chol, 4% PI, and 1% PIP_2_. Physiological ionic strength with 1 mM MgCl_2_/3 mM ATP was used.

**Supplementary Figure 4 No effect of CaM on Ca^2+^-dependent SV fusion. (a)** SV fusion with PM-liposomes using a lipid-mixing assay as in Fig. 1b. Addition of 100 µM free Ca^2+^ accelerated SV fusion, but 5 µM CaM had no effect on Ca^2+^-dependent SV fusion. Physiological ionic strength with 1 mM MgCl_2_/3 mM ATP was used.

**Supplementary Figure 5 Mathematical modelling of the interaction between charged membranes.** (A) Schematic diagram of two lipid vesicles in aqueous solution at separation distance *R*. (B) Schematic representation of two membranes separated by an aqueous phase at a distance *R*. *h*, hydrophilic polar layers.

**Supplementary Figure 6 Mathematical modelling of membrane attraction by the lipid tail hydrophobicity.** (A) van der Waals, electrostatic, hydration, and the total energy between two membranes when the hydrophobicity index *p* = 0.6. (B) Total interaction energy at the hydrophobicity index *p* = 0.0, 0.2, 0.4, 0.6, 0.8, and 1.0.

## Online Methods

### Materials

ATP disodium salt was from Sigma-Aldrich (Cat. A2383). Antibody against to VAMP-2 (synaptobrevin-2) was purchased from Synaptic Systems (clone number 69.1), Göttingen, Germany. Alexa Fluor 488 C5 maleimide (Cat. A10254), BODIPY 493⁄503 Succinimidyl Ester (Cat. D2191), and BODIPY TR-X Succinimidyl Ester (Cat. D6116) were purchased from Thermo Fisher Scientific. The MARCKS effector domain (residues 151–175 of bovine MARCKS, termed MARCKS ED) consisting of KKKKKRFSFKKSFKLSGFSFKKNKK was synthesized by GenScript (Piscataway, NJ). Recombinant human PKC-epsilon (Cat. SRP5067), recombinant bovine CaM (Cat. C4874), PMA (Cat. P8139), and GF109203X (Cat. B6292) were purchased from Sigma-Aldrich. All lipids were purchased from Avanti Polar lipids except Oregon Green 488 1,2-Dihexadecanoyl-*sn*-Glycero-3-Phosphoethanolamine (Oregon Green 488 DHPE) (Cat. O12650, Thermo Fisher Scientific).

### Purification of large dense-core vesicles (LDCVs) and synaptic vesicles (SVs)

LDCVs, also known as chromaffin granules, were purified from bovine adrenal medullae by using continuous sucrose gradient and resuspended with fusion buffer containing 120 mM K-glutamate, 20 mM K-acetate, and 20 mM HEPES.KOH, pH 7.4, as described previously^44^. Briefly, fresh bovine adrenal glands were obtained from a local slaughterhouse. The cortex and fat were removed, then the medullae were minced with scissors in 300 mM sucrose buffer (300 mM sucrose, 20 mM HEPES, pH 7.4 adjusted with KOH), and homogenized using a homogenizer. After centrifugation at 1,000g for 15 min at 4 °C, the pellet containing nuclei and cell debris (P1) was discarded. The supernatant (S1) was further centrifuged (12,000g, 15 min, 4 °C), then subjected to an additional cycle of resuspension and centrifugation as a washing step. The resulting pellet (P2, crude LDCV fraction) was resuspended in 300 mM sucrose buffer and loaded on top of a continuous sucrose gradient (from 300 mM to 1.9 M) to remove other contaminants including mitochondria. LDCVs were collected from the pellet after centrifugation at 110,000g for 60 min in a Beckman SW 41 Ti rotor, then resuspended with the buffer (120 mM K-glutamate, 20 mM K-acetate, 20 mM HEPES.KOH, pH 7.4).

SV from mouse brains were purified as described elsewhere^68^. Briefly, the brains were homogenized in homogenization buffer supplemented with protease inhibitors, using a glass-Teflon homogenizer. The homogenate was centrifuged for 10 min at 1,000g, and the resulting supernatant was further centrifuged (15 min, 15,000g, 4 °C). The synaptosome pellet was lysed by adding ice-cold water, followed by centrifugation (25 min, 48,000g, at 4 °C). The resulting supernatant was overlaid onto a 0.7 M sucrose cushion and centrifuged for 1 h at 133,000g. The pellet was resuspended in fusion buffer (120 mM K-glutamate, 20 mM K-acetate, 20 mM HEPES.KOH, pH 7.4).

### Protein purification

All SNARE and synaptotagmin-1 constructs based on *Rattus norvegicus* sequences were expressed in *E. coli* strain BL21 (DE3) and purified by Ni^2+^-NTA affinity chromatography followed by ion-exchange chromatography as described elsewhere^17, 18^. The stabilized Q-SNARE complex was composed of syntaxin-1A (aa 183–288) and SNAP-25A (no cysteine, cysteines replaced by alanines) in a 1:1 molar ratio by the C-terminal VAMP-2 fragment (aa 49–96), and was purified as described earlier^45^. The soluble stabilized Q-SNARE complex with syntaxin-1A (aa 183-262) was purified as described earlier^45^. The binary Q-SNARE complex containing the full-length syntaxin-1A (1-288) and SNAP-25A (no cysteine, cysteines replaced by alanines) was expressed using co-transformation^17^. The full-length VAMP-2, soluble cytoplasmic region of VAMP-2 (VAMP-2_1-96_), full-length synaptotagmin-1, and the C2AB domain of synaptotagmin-1 (aa 97-421) were purified as described previously^69^. The stabilized Q-SNARE complex and the syntaxin-1A/SNAP-25A binary SNARE complex were purified by Ni^2+^-NTA affinity chromatography followed by ion-exchange chromatography on a Mono Q column (GE Healthcare, Piscataway, NJ) in the presence of 50 mM n-octyl-β-D-glucoside (OG)^17^. The point-mutated C2AB domain (S342C)(C2AB-Alexa 488)^69^ and VAMP-2_49-96_ (T79C)^17, 45^ in the stabilized Q-SNARE complex were labelled with Alexa Fluor 488 C5 maleimide. MARCKS ED was labeled at the N-terminus with BODIPY 493⁄503 Succinimidyl Ester or BODIPY TR-X Succinimidyl Ester. Syntaxin-1A (1-288) was labelled with Alexa Fluor 594 C5 maleimide at T197C^18^. Protein structures were visualized with PyMOL; PDB 1BYN for the C2A domain, 1K5W for the C2B domain, 3IPD for the SNARE complex, 1CLL for CaM.

### Lipid composition of liposomes

All lipids were from Avanti Polar lipids, unless stated otherwise; Oregon Green 488 DHPE was from Thermo Fisher Scientific. Lipid composition (molar percentages) of PM-liposomes containing the Q-SNARE complex: 45% PC (L-α-phosphatidylcholine, Cat. 840055), 15% PE (L-α-phosphatidylethanolamine, Cat. 840026), 10% PS (L-α-phosphatidylserine, Cat. 840032), 25% Chol (cholesterol, Cat. 700000), 4% PI (L-α-phosphatidylinositol, Cat. 840042), and 1% PIP_2_ (Cat. 840046). When PIP_2_ was excluded (0% PIP_2_) or changed, PI contents were accordingly adjusted. In case of removing PS/PIP_2_, PC contents were accordingly adjusted. To measure PS-dose dependence on vesicle fusion, lipid composition of PM-liposomes was 20% PC, 15% PE (labelled PE included), 40% PS, and 25% Chol. When PS contents were changed, PC contents were accordingly adjusted. VAMP-2/synaptotagmin-1-containing V-liposomes were composed of 55% PC, 20% PE, 15% PS, and 10% Chol.

For vesicle fusion lipid-mixing assays, 1.5% 1,2-dioleoyl-*sn*-glycero-3-phosphoethanolamine-N-(7-nitrobenz-2-oxa-1,3-diazol-4-yl) (NBD-PE, Cat. 810145) as a donor and 1.5% 1,2-dioleoyl-*sn*-glycero-3-phosphoethanolamine-N-lissamine rhodamine B sulfonyl ammonium salt (Rhodamine-PE, Cat. 810150) as an acceptor dye were incorporated in PM-liposomes (accordingly 12% unlabelled PE).

For FRET measurement using the C2AB domain labeled with Alexa 488, 1.5% Rhodamine-DOPE was included in liposomes as an acceptor dye; lipid composition of liposomes: protein-free, 45% PC, 13.5% PE, 1.5% Rho-PE, 10% PS, 25% Chol, 4% PI, and 1% PIP_2_.

For FRET measurement using MARCKS ED labeled with BODIPY TR, 1.5% Oregon Green 488 DHPE (Cat. O12650, Thermo Fisher Scientific) was included in liposomes as a donor dye; lipid composition of liposomes: protein-free, 45% PC, 13.5% PE, 1.5% Oregon Green 488-DHPE, 10% PS, 25% Chol, 4% PI, and 1% PIP_2_. If PS/PIP_2_ were removed, PC contents were accordingly adjusted.

For anisotropy measurement, lipid composition of liposomes (no fluorescent-labeled lipids included): 45% PC, 15% PE, 10% PS, 25% Chol, 4% PI, and 1% PIP_2_. In **Supplementary Figure 2a**, MARCKS ED was labeled with BODIPY 493⁄503 and lipid composition of liposomes was 75% PC, 25% Chol vs. 74% PC, 25% Chol, 1% PIP_2_.

### Preparation of proteoliposomes

Incorporation of the stabilized or binary Q-SNARE complex into large unilamellar vesicles (LUVs) was achieved by OG-mediated reconstitution, called the direct method, i.e. incorporation of proteins into preformed liposomes^17, 18^. LUVs prepared by the direct method were used, unless stated otherwise. Briefly, lipids dissolved in chloroform were mixed according to lipid composition. The solvent was removed using a rotary evaporator (which generated lipid film on a glass flask), then lipids were resuspended in 0.5 mL buffer containing 150 mM KCl and 20 mM HEPES/KOH pH 7.4. After sonication on ice, multilamellar vesicles were extruded using polycarbonate membranes of pore size 100 nm (Avanti Polar lipids) to give uniformly-distributed LUVs with average diameter 110 nm^20^. After the preformed LUVs had been prepared, SNARE proteins (for PM-liposomes) or the full-length VAMP-2/synaptotagmin-1 (for V-liposomes) were incorporated into liposomes by using OG, a mild non-ionic detergent, then OG was removed by dialysis overnight in 1 L buffer containing 150 mM KCl and 20 mM HEPES/KOH pH 7.4 together with 2 g SM-2 adsorbent beads.

Small unilamellar vesicles (SUVs, ∼50 nm in diameter, **Supplementary Fig. 1d**) were produced by the standard method, i.e. co-solubilizing proteins and lipids by using a strong detergent. Briefly, lipid film was dissolved in 25 µL buffer containing 150 mM KCl, 20 mM HEPES/KOH pH 7.4, and 5% sodium cholate. In parallel, SNARE proteins (for PM-liposomes) or the full-length VAMP-2/synaptotagmin-1 (for V-liposomes) were resuspended in 75 µL buffer containing 150 mM KCl, 20 mM HEPES/KOH pH 7.4, and 1.5% sodium cholate. Lipids and proteins were combined (100 µL in total), then a size-exclusion column was used to remove detergent (Sephadex G50 in 150 mM KCl and 20 mM HEPES, pH 7.4). SUVs were collected from the size-exclusion column as eluted (∼ 400 µL); liposomes were easily detected visually due to fluorescent-labelled lipids. The size distribution of proteoliposomes was determined using dynamic light scattering (DLS)^20, 44^. Protein to lipid ratio in proteoliposomes was 1:500 (n/n).

### Vesicle fusion assay

A FRET-based lipid-mixing assay was performed to monitor native vesicle fusion *in vitro*^17, 18, 20, 43, 44^. LDCV or SV fusion reactions were performed at 37°C in 1 mL fusion buffer containing 120 mM K-glutamate, 20 mM K-acetate, 20 mM HEPES-KOH (pH 7.4), 1 mM MgCl_2_, and 3 mM ATP. ATP should be made freshly before experiments, because it is easily destroyed by freezing and thawing. Free Ca^2+^ concentration in the presence of ATP and Mg^2+^ was calibrated using the MaxChelator simulation program. The fluorescence dequenching signal was measured using Fluoromax (Horiba Jobin Yvon) with wavelengths of 460 nm for excitation (Ex) and 538 nm for emission (Em). Fluorescence values were normalized as a percentage of maximum donor fluorescence (total fluorescence) after addition of 0.1% Triton X-100 at the end of experiments.

### Fluorescence anisotropy measurements

Anisotropy measurement^18, 42^ was carried out at 37°C in 1 mL buffer containing 120 mM K-glutamate, 20 mM K-acetate, and 20 mM HEPES-KOH (pH 7.4), 1 mM MgCl_2_, and 3 mM ATP. Anisotropy (*r*) was calculated as *r* = (I_VV_ − G × I_VH_)/(I_VV_ + 2 ×G × I_VH_), where I_VV_ denotes the fluorescence intensity with vertically polarized excitation and vertical polarization on the detected emission, and I_VH_ denotes the fluorescence intensity when using a vertical polarizer on the excitation and horizontal polarizer on the emission. G is a grating factor used as a correction for the instrument’s differential transmission of the two orthogonal vector orientations. Lipid composition of PM-liposomes (protein-free) was identical to those used in a fusion assay except labelled PE (45% PC, 15% PE, 10% PS, 25% Chol, 4% PI, and 1% PIP_2_). Anisotropy (a.u.) was presented as A/A_0_, where A_0_ is the initial value. The C2AB domain labeled with Alexa Fluor 488, Ex/Em = 495/520 nm; MARCKS ED labeled with BODIPY TR, Ex/Em = 590/620 nm; MARCKS ED labeled with BODIPY 493⁄503, Ex/Em = 493/503 nm.

### Fluorescence resonance energy transfer (FRET)

The C2AB domain labeled with Alexa 488 (a donor dye) was incubated with liposomes that include 1.5% Rhodamine-DOPE (an acceptor dye); donor fluorescence signal was measured with wavelengths of 488 nm for excitation and 516 nm for emission^20^. MARCKS ED labeled with BODIPY TR (an acceptor dye) was incubated with liposomes that incorporate 1.5% Oregon Green 488 DHPE (a donor dye); donor fluorescence signal was measured with wavelengths of 500 nm for excitation and 540 nm for emission. Unless otherwise stated, liposomes were LUVs prepared by the direct method. Donor fluorescence signal was measured at 37°C using Fluoromax (Horiba Jobin Yvon) in 1 mL buffer containing 120 mM K-glutamate, 20 mM K-acetate, 20 mM HEPES-KOH (pH 7.4), 1 mM MgCl_2_, and 3 mM ATP. FRET was normalized as F/F_0_, where F_0_ represents the initial value of the donor fluorescence intensity.

### Ternary SNARE complex formation assay

Tetanus neurotoxin (TeNT) degrades free VAMP-2 whereas VAMP-2, engaged in the ternary SNARE complex, is resistant to TeNT^17^. After incubation of LDCVs with PM-liposomes that contain the stabilized Q-SNARE complex for 20 min at 37 °C, the sample was subjected to TeNT treatment (200 nM, 30 min, 37°C), then boiled for 5 min at 95°C and analysed by immunoblotting with antibody against VAMP-2 (clone number 69.1, Synaptic Systems, Göttingen, Germany).

### Preparation of bovine chromaffin cells

Chromaffin cells were isolated from the bovine adrenal gland medulla by two-step collagenase digestion as previously described^70, 71^. The cells were grown on poly-D-lysine-coated glass coverslips in Dulbecco’s modified Eagle medium/F-12 (Cat. 11320033, Gibco™) containing 10% fetal bovine serum (Cat. SH3007103, Cytiva) and 1% antibiotics (Cat. 10378016, Gibco™).

### Amperometric measurement

Recordings of LDCV exocytosis from chromaffin cells were performed as described previously^70^. Chromaffin cells were buffered with amine-free solution containing 137. 5 mM NaCl, 2.5 mM KCl, 2 mM CaCl_2_, 1 mM MgCl_2_, 10 mM D-glucose, and 10 mM HEPES-NaOH (pH 7.3). Carbon-fiber electrodes were fabricated with 8 μm diameter carbon fibers and back-filled with 3 M KCl. The amperometric current was generated by oxidation of catecholamine, and was measured using an axopatch 200B amplifier (Axon Instruments Inc., CA), which was operated in a voltage-clamp mode at a holding potential of + 650 mV. Amperometric signals were low-pass filtered at 1 kHz and sampled at 500 Hz. For data acquisition and analysis, pCLAMP 11 software (Axon Instruments) was used.

### Transmission electron microscopy (TEM)

As described previously^20^, 5 µL of samples were deposited on carbon-coated 400-mesh copper grids (CF400-CU, Electron Microscopy Sciences). Grids were stained with uranyl acetate for negative staining and embedded in uranyl acetate. Vesicles were visualized at 80 kV in Talos F200C Transmission Electron Microscope (Thermo Fisher Scientific). The images were acquired using bottom-mounted CETA camera.

### Mathematical modelling

For the interaction between charged phospholipid vesicles, we used the modified Derjaguin-Landau-Verwey-Overbeek (DLVO) theory suggested by Ohki and Oshima (**Supplementary Fig. 5**).^52^ When two membranes interact at a close distance, the hydration energy contribution to the total energy becomes dominant, and the total energy can be expressed as the sum of the van der Waals energy, the electrostatic energy, and the hydration energy.

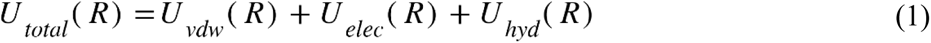

The van der Waals energy can be expressed as equation (2).

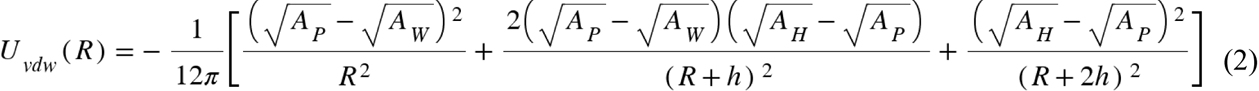

*A*_W_, the Hamaker constant of water phase (3.0 × 10^−13^ erg); *A*_H,_ the Hamaker constant of hydrocarbon (6.5 × 10^−13^ erg); *A*_P_, the Hamaker constant of the polar phase, the mixture of the water and hydrocarbon phases is expressed as

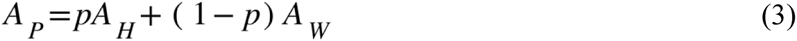

where *p* is the degree of hydrophobicity of the surface layer (0 ≤ *p* ≤ 1). The electrostatic interaction energy can be expressed as equation (4).

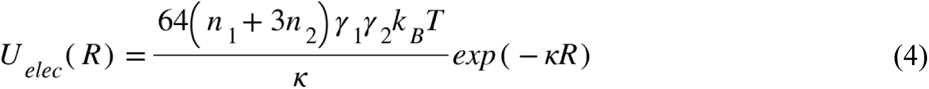

where *n*_1_ and *n*_2_ are bulk concentration of 1-1 and 2-1 electrolytes, κ is the Debye length, 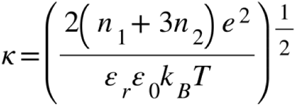, and *y*_0j_ is the scaled surface potential 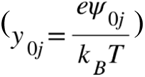 of the membrane surface potential, *Ψ*_0j_. γ and η is expressed as follows.

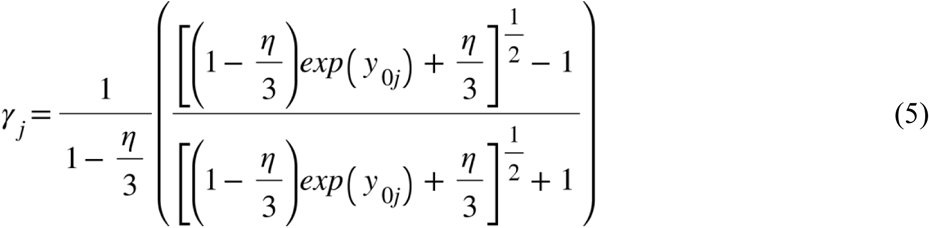

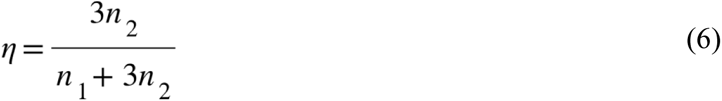

The detailed derivation of the equations and their explanations are shown in the works of Oshima and Ohki.^52, 72^ The hydration energy of the contacting membranes is shown in equation (7).

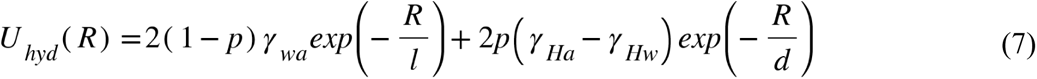

*γ*_wa_, *γ*_Ha_, and *γ*_Hw_ is the interfacial surface tension of water/air (∼72 dyne/cm), hydrocarbon/air (∼20 dyne/cm), and hydrocarbon/water (∼50 dyne/cm) phases. The short (*l*) and long decay (*d*) constants of the hydration energy were taken as 2 and 10 Å based on the previous works by Oshima and Ohki.^52^ When the surface potential of the synaptic vesicle *Ψ*_0_ = –20 mV^73^ and the surface potential of the phospholipid with 5 – 20 % PIP2 is in the range of –20 ∼ –80 mV,^74^ the total interaction energy between two parallel phospholipid membranes is shown in **Supplementary Fig. 6**.

### Statistical analysis

Data analysis was performed using OriginPro 2019 software (OriginLab Corporation, Northampton, MA, USA) and GraphPad Prism 9 (GraphPad Software, San Diego, CA, USA). Data are means ± standard deviation (SD) or standard error of mean (SEM). One-way ANOVA test with Bonferroni correction or post-hoc Turkey was used to determine any statistically significant differences among three or more independent groups. Probabilities *p* < 0.05 were considered significant. Dose-response curves were fitted using four-parameter logistic equations (GraphPad Prism) to calculate IC_50_.

