## Supplementary figures for "PIP_2_ electrostatically triggers vesicle fusion: arresting full SNARE assembly and vesicle fusion by PIP_2_-masking"

### Supplementary Figure 1

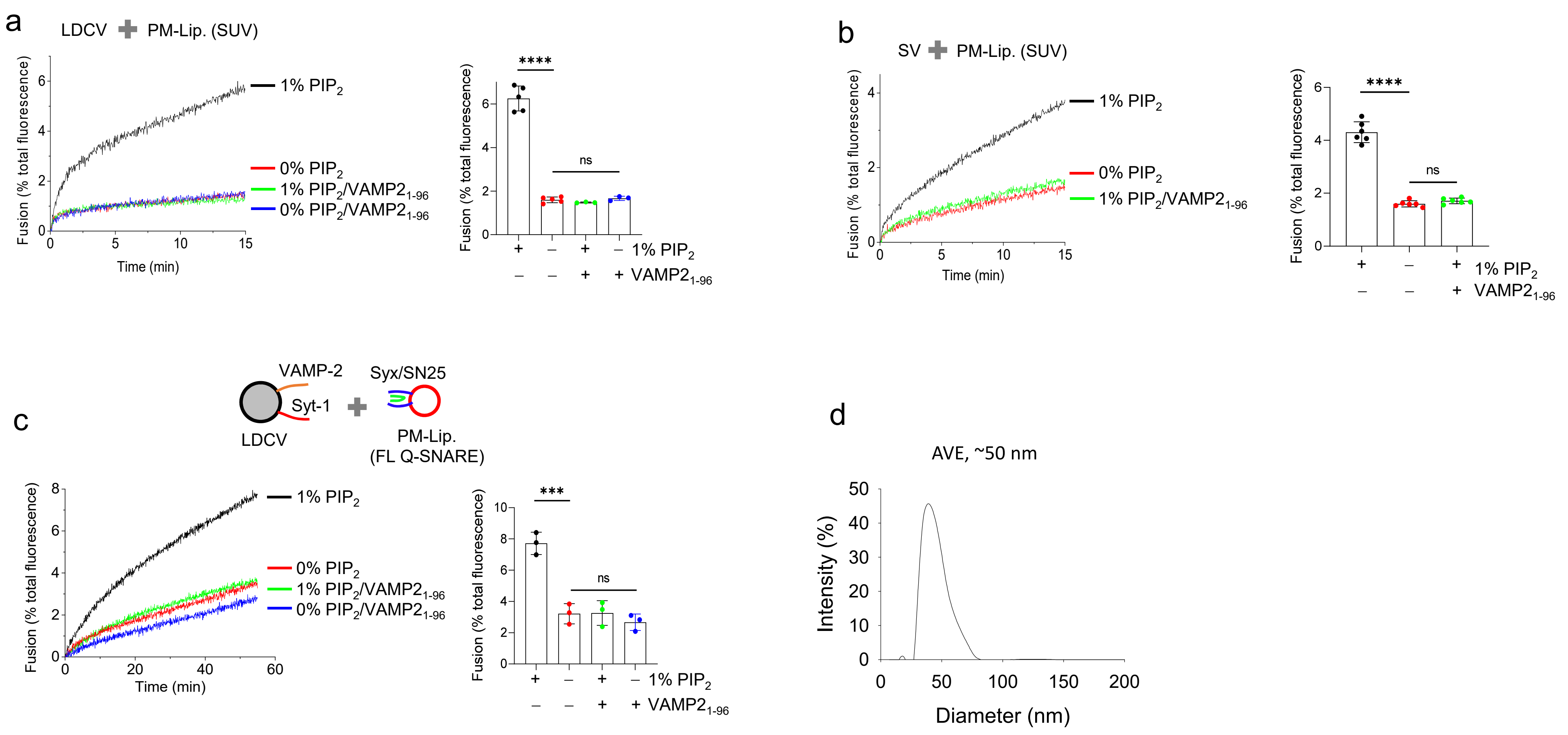

### Supplementary Figure 2

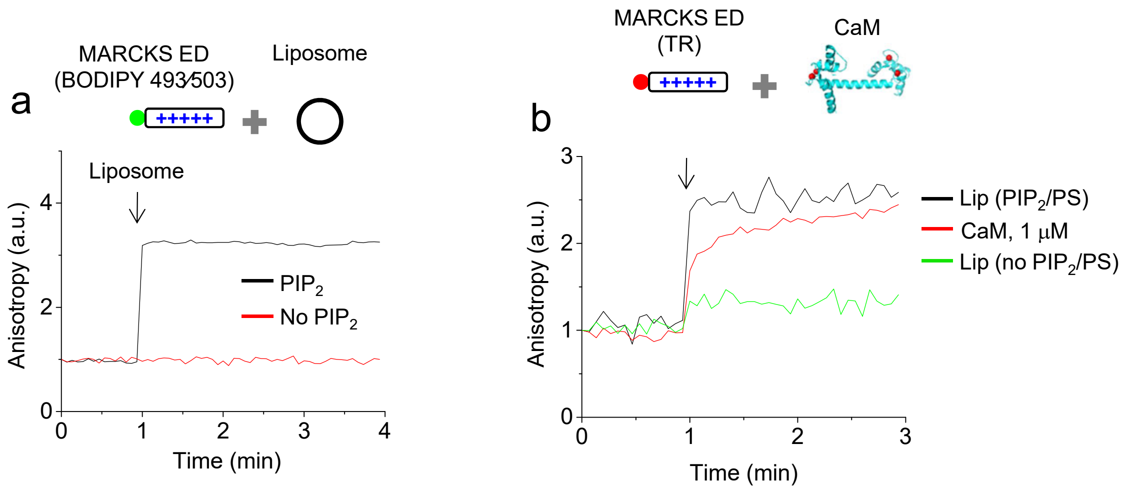

### Supplementary Figure 3

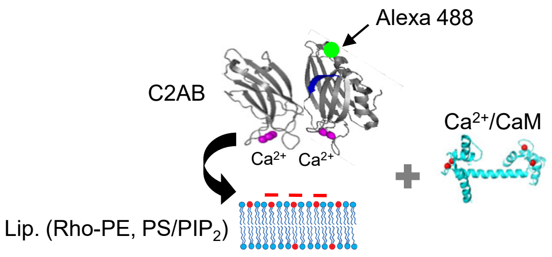


a

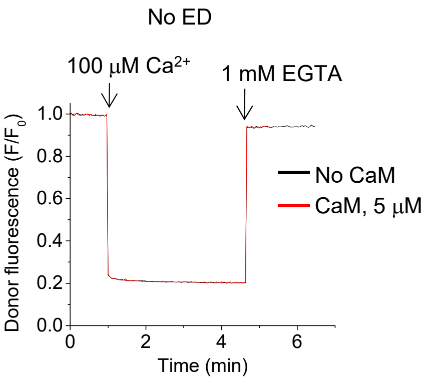

### Supplementary Figure 4

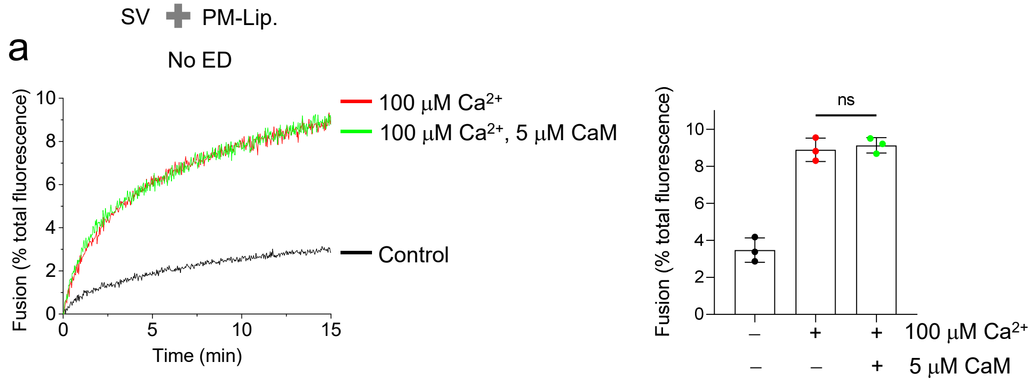

Supplementary Figure 5

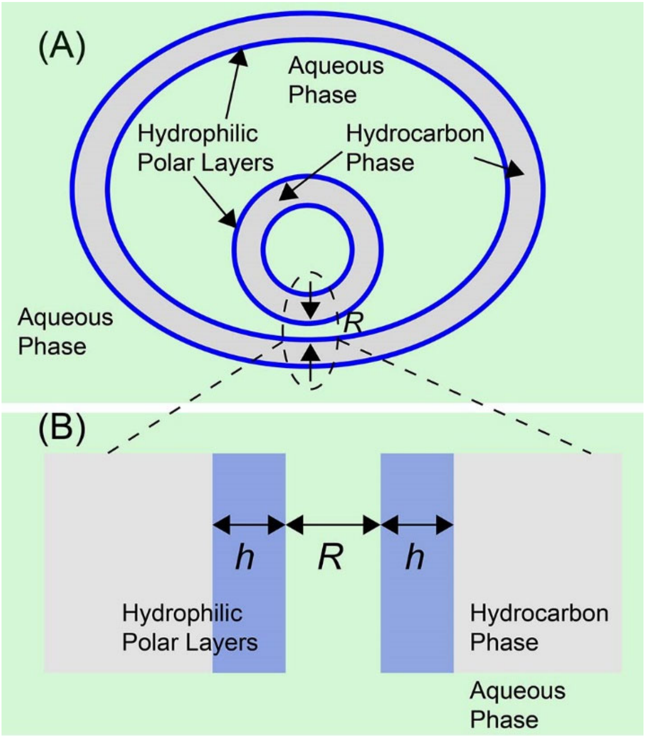

Supplementary Figure 6

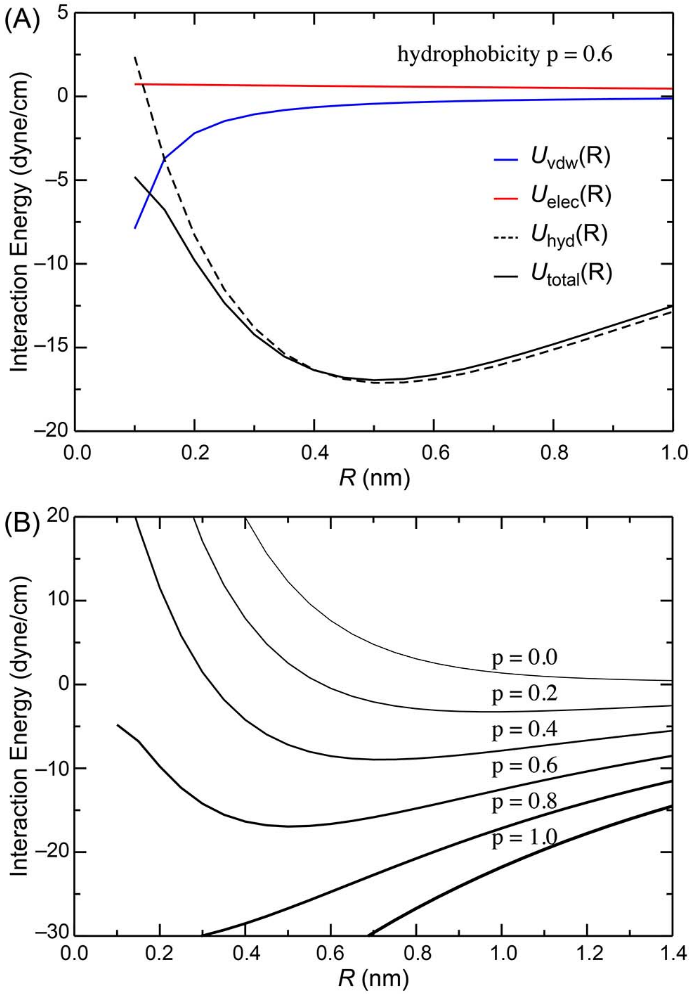
